# Sensory event-related potential morphology predicts age in premature infants

**DOI:** 10.1101/2023.07.21.549656

**Authors:** Coen S. Zandvoort, Marianne van der Vaart, Shellie Robinson, Fatima Usman, Gabriela Schmidt Mellado, Ria Evans Fry, Alan Worley, Eleri Adams, Rebeccah Slater, Luke Baxter, Maarten de Vos, Caroline Hartley

## Abstract

Preterm infants undergo substantial neurosensory development in the first weeks after birth. Infants born prematurely are more likely to have long-term adverse neurological outcomes and early detection of abnormal brain development is essential for timely interventions. We investigated whether sensory-evoked cortical potentials could be used to accurately estimate the age of an infant. Such a model could be used to identify infants who deviate from normal neurodevelopment by comparing the brain age to the infant’s postmenstrual age (PMA). Infants aged between 28- and 40-weeks PMA from a training and test sample (consisting of 101 and 65 recording sessions in 82 and 14 infants, respectively) received trains of approximately 10 visual and 10 tactile stimuli (interstimulus interval approximately 10 seconds). PMA could be predicted accurately from the magnitude of the evoked responses (training set mean absolute error (MAE and 95% confidence intervals): 1.41 [1.14; 1.74] weeks, *p* = 0.0001; test set MAE: 1.55 [1.21; 1.95] weeks, *p* = 0.0002. Moreover, we show with two examples that brain age, and the deviations between brain age and PMA, may be biologically and clinically meaningful. By firstly demonstrating that brain age is correlated with a measure known to relate to maturity of the nervous system (based on animal and human literature, the magnitude of reflex withdrawal is used) and secondly by linking brain age to long-term neurological outcomes, we show that brain age deviations are related to biologically meaningful individual differences in the rate of functional nervous system maturation rather than noise generated by the model. In summary, we demonstrate that sensory-evoked potentials are predictive of age in premature infants. It takes less than 5 minutes to collect the stimulus electroencephalographic data required for our model, hence, increasing its potential utility in the busy neonatal care unit. This model could be used to detect abnormal development of infant’s response to sensory stimuli in their environment and may be predictive of later life abnormal neurodevelopmental outcome.

## Introduction

Premature and hospitalised infants are at increased risk of adverse neurodevelopmental outcomes compared with healthy term-born infants (Blencowe et al., 2013). The neurosensory system of premature infants undergoes rapid structural and functional development (Kostović et al., 2014; Niemarkt et al., 2011), with functional changes apparent in electroencephalographic (EEG) recordings (André et al., 2010). Sensory-evoked potentials provide information about the integrity of the sensory nervous system and may be predictive of neurological outcomes (Leikos et al., 2020; Majnemer and Rosenblatt, 1996; Pike and Marlow, 2000; Taylor et al., 1996). A variety of neural impairments associated with atypical development of the somatosensory and visual systems have been described, affecting both the morphology and latency of evoked potentials (De Vries et al., 1990; de Zegher et al., 1992; Häkkinen et al., 1987; McCulloch et al., 1991; Taylor and McCulloch, 1992; Whyte et al., 1987). Generally, sensory stimuli evoke slow-wave responses in young premature babies (Khazipov et al., 2004), whereas evoked brain activity with high-frequency waveforms are observed in older infants (André et al., 2010; Niemarkt et al., 2011).

Machine learning approaches can be used to accurately predict the post-menstrual age (PMA) of preterm infants from EEG (Ansari et al., 2023; Lavanga et al., 2018; O’Toole et al., 2016; Pillay et al., 2020; Stevenson et al., 2017), diffusion magnetic resonance imaging (MRI) (Brown et al., 2017; Kawahara et al., 2017) and structural MRI (Liu et al., 2021). These models may facilitate the early identification of infants with abnormal neurodevelopment, reducing the need for visual inspection of the EEG/MRI, which is subjective, requires trained clinical staff, and is time-consuming. This so-called brain age can be seen as a maturation index of the neural system which is unlikely to reflect chronological age, which can be viewed as a continuous “ticking clock” (Salih et al., 2023). Previous EEG brain age models have focused on continuous ongoing EEG activity (i.e., non-evoked brain activity) (Ansari et al., 2023; O’Toole et al., 2016; Pillay et al., 2020; Stevenson et al., 2017). An alternative may be to construct models capturing evoked responses, giving specific information about the maturity of sensory processing. We hypothesised that sensory-evoked responses will be predictive of age and that development of brain age models which use sensory-evoked potentials may specifically provide insight into neurosensory brain functioning in premature infants.

Here, we aimed to assess whether sensory-evoked responses could be used to predict PMA in infants, focusing on visual and tactile stimuli as these are easy to perform in infants and elicit clear evoked potentials requiring only a small number of trials. To facilitate the development of a sensory-evoked brain age model, we utilise stimulus-specific neurodynamic response functions (NRF), which, akin to haemodynamic response functions used in functional MRI (fMRI) (Arichi et al., 2012; Henson and Friston, 2007), represent the characteristic waveforms evoked by the stimuli. Identifying NRFs provides a consistent reproducible approach to compare infants across research studies (Hartley et al., 2017) and is likely a useful candidate feature for predicting age (Green et al., 2019; Hartley et al., 2016; Schmidt Mellado et al., 2022; van der Vaart et al., 2022). NRFs have been previously developed for responses to visual (Schmidt Mellado et al., 2022), tactile (Schmidt Mellado et al., 2022), and noxious (Hartley et al., 2017) stimuli in term infants. These term-derived brain responses show that sensory-evoked potentials change with age in premature infants (Fabrizi et al., 2011; Hartley et al., 2016; Schmidt Mellado et al., 2022; van der Vaart et al., 2022), are sensitive to stimulus intensity (Hartley et al., 2015), and may be modulated by interventions (Cobo et al., 2021; Gursul et al., 2018; Hartley et al., 2017); however, deriving NRFs from preterm infants across development will be better able to predict brain age.

In this study, we first identified NRFs of visual- and tactile-evoked brain activity in infants between 28-40 weeks PMA. Next, we quantified age-dependent relationships for each of the NRFs and trained a machine learning model that accurately predicted brain age using these NRFs. In an independent sample of preterm infants, we tested the NRFs and age prediction model. Finally, in two examples we explored whether the infants’ brain ages are meaningful. Firstly, we tested if the magnitude of reflex withdrawal is correlated with infant brain age, suggesting its biological significance. Secondly, we related longitudinal brain age trajectories to long-term outcomes and expected that infants with below-average Bayley-III outcomes would have greater differences between their brain age and PMA (i.e., greater brain age gap) and different brain development trajectories when compared with infants with average Bayley-III outcomes. This would suggest that brain age trajectory (identified using the sensory evoked model presented here) may be clinically meaningful and predictive of later life outcome.

## Results

### Stimulus-evoked potentials change with post-menstrual age

Stimulus-evoked EEG responses to visual and tactile stimuli could be observed between 28 and 40 weeks PMA with distinct morphological changes across this age range (Figure 1, training set and Figure S1, test set). In response to the visual stimulus, a low frequency waveform with negative polarity was observed at the Oz channel in the youngest infants, which disappeared with increasing age (first row of Figure 1). A higher frequency potential was present across all ages, with apparent shift in latency and morphology. Following tactile stimulation, the very youngest infants also displayed a slow-wave response whereas older infants displayed a clear negative peak at ∼0.16 s post-stimulus (second row of Figure 1). The test set demonstrated waveforms of similar morphology to the training set across the age range studied (Figure S1). Note that stimulus responses and age-prediction models were first derived in a training set and then validated in an independent test set; however, for ease of comparison, data in the test set is presented together with the training set throughout the remaining of the results.

**Figure 1.**
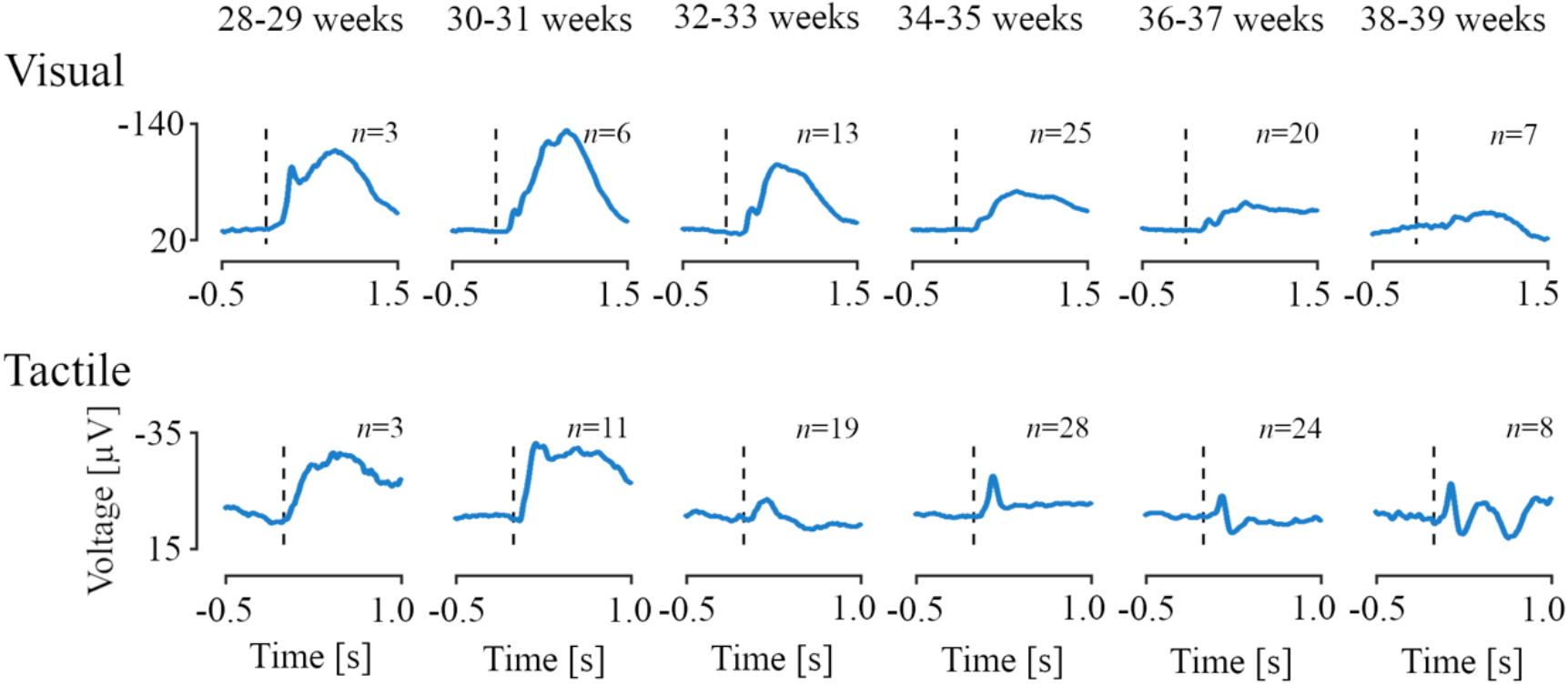
Stimulus-evoked electroencephalographic potentials according to infant age. Age-dependent evoked potentials for two-weeks intervals between 28 to 40 weeks of post-menstrual age for the visual and tactile stimuli at channels Oz and Cz, respectively. For the test data set, evoked responses are comparable (see Figure S1). Woody filtering aligned the responses to their age-weighted averages. Vertical dashed lines correspond to the stimulus onset. Number of infants indicated by n.

### Stimulus-evoked potentials can be characterised using neurodynamic response functions

We used a data-driven approach to identify the characteristic waveforms (the NRFs) from the visual- and tactile-evoked potentials of the training set. Visual-evoked activity at Oz occurred between 0.23 and 1.0 s post-stimulus (permutation testing, *p* = 0.001, Figure S2). Tactile evoked activity at channel Cz occurred between 0.09 to 0.23 s post-stimulation (*p* = 0.041, Figure S2). Four NRFs were identified in response to the visual stimulus and two NRFs in response to the tactile stimulus (Figure 2a and Table S1). In the test sample, the magnitudes of all NRFs were significantly different between the stimulus-evoked activity and resting state, demonstrating the reproducibility of these response functions in an independent dataset (Figure S3 and Table S1). NRFs 1 and 2 in response to the visual stimulus consist of low-frequency waves. NRF 1 also has a superimposed higher frequency waveform at ∼0.27 s (Figure 2a). Visual NRFs 3 and 4 are higher frequency components with rapid negative-positive polarity changes from ∼0.25 up to 1 s (Figure 2a). The magnitude of the visual-evoked brain activity for the NRFs change with age (particularly in the training set), indicating that these responses may be useful features for a brain age prediction model (linear regressions were used as a guide, Figure 2b-c and Table S2).

**Figure 2.**
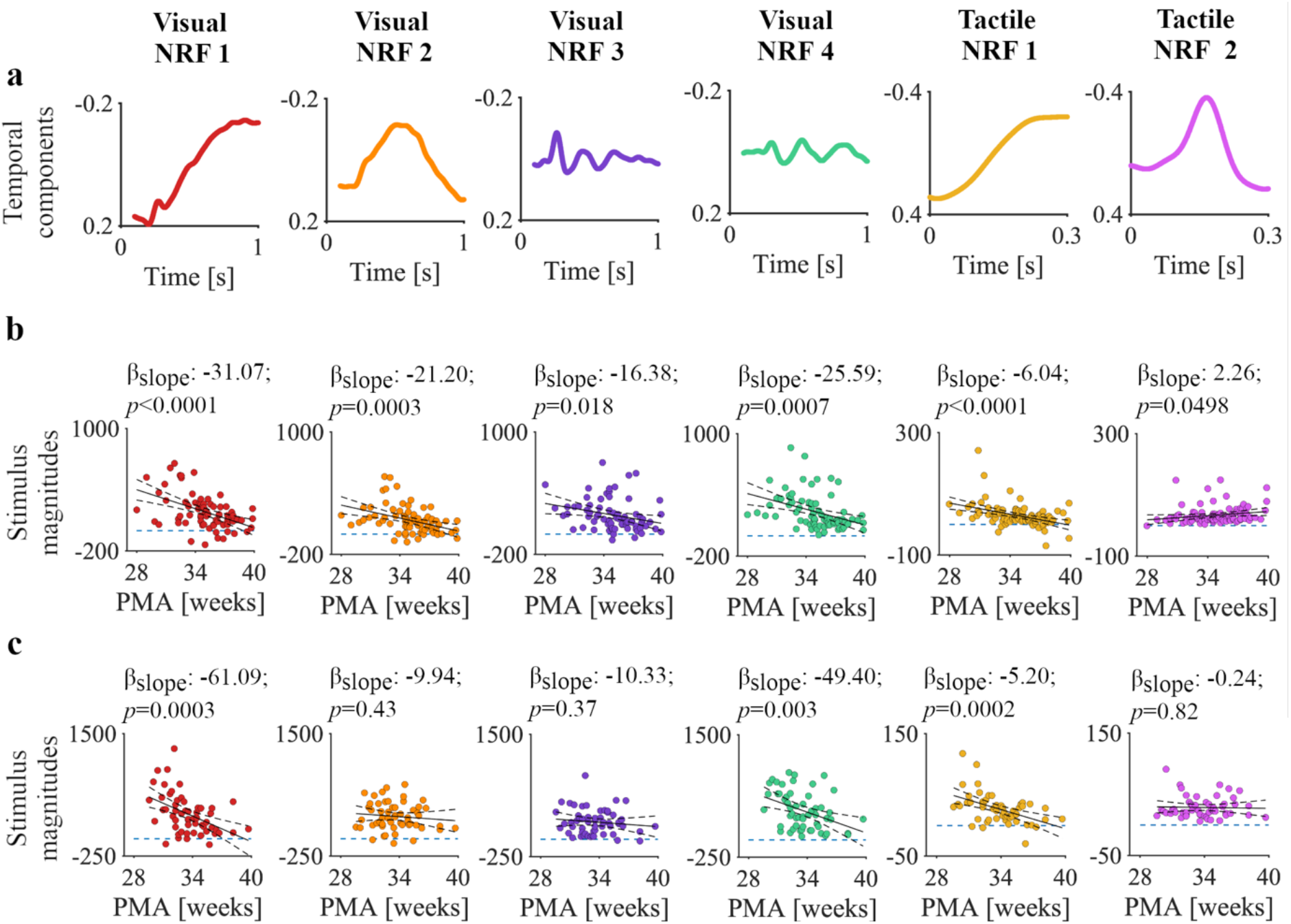
Waveforms of the neurodynamic response functions (NRFs) and magnitude changes with post-menstrual age (PMA). **a)** NRFs as a function of time identified from the training sample. Six (four visual and two tactile) principal components revealed statistically significant mean differences in NRF magnitudes between stimulus responses and resting state activity. **b)** The relationships between PMA and NRF magnitudes for each recording in the training sample (n = 74 and n = 93 for the visual and tactile responses, respectively). Continuous and dashed black graphs are the fitted means and 95%-confidence intervals of the generalised linear models. Dashed horizontal blue lines mark a magnitude of 0. **c)** The relationships between PMA and NRF magnitudes for each recording in the test sample (visual and tactile responses comprised 55 and 53 recordings, respectively).

Tactile NRF 1 consists of a slow-wave component (Figure 2a), of which the magnitudes significantly decreased with PMA in the training and test samples (Figure 2b-c and Table S2). Tactile NRF 2 is a higher frequency component with a negative deflection at ∼0.17 s (Figure 2a); the stimulus response is present at all PMAs in the training and test samples, except for one infant at 28 weeks in the training sample (Figure 2b-c and Table S2). Visual inspection of the NRFs projected on age-specific averages and recording-averaged responses (Figures S4-S11) demonstrated a good fit within individual subjects and age-dependent changes in responses. To summarise, the characteristic waveforms from visual and tactile responses show changes with PMA in both training and test sets.

### Visual and tactile-evoked responses are predictive of the age of the infant

Using the NRF magnitudes of the stimulus responses, we used support vector regression to build a model which could accurately predict infant age (Figure 3a-b, training sample leave-one-infant-out cross-validation, MAE = 1.41 weeks with a 95% confidence interval of [1.14; 1.74] weeks, *p* = 0.0001). In the independent test sample, this model accurately predicted the age of the infants (Figure 3c-d, MAE = 1.55 weeks with 95% CI at [1.21; 1.95] weeks, *p* = 0.0002). Models trained on the responses to either the visual or tactile stimuli only did not perform significantly better than the null models in the test set; Figures S12-S13).

**Figure 3.**
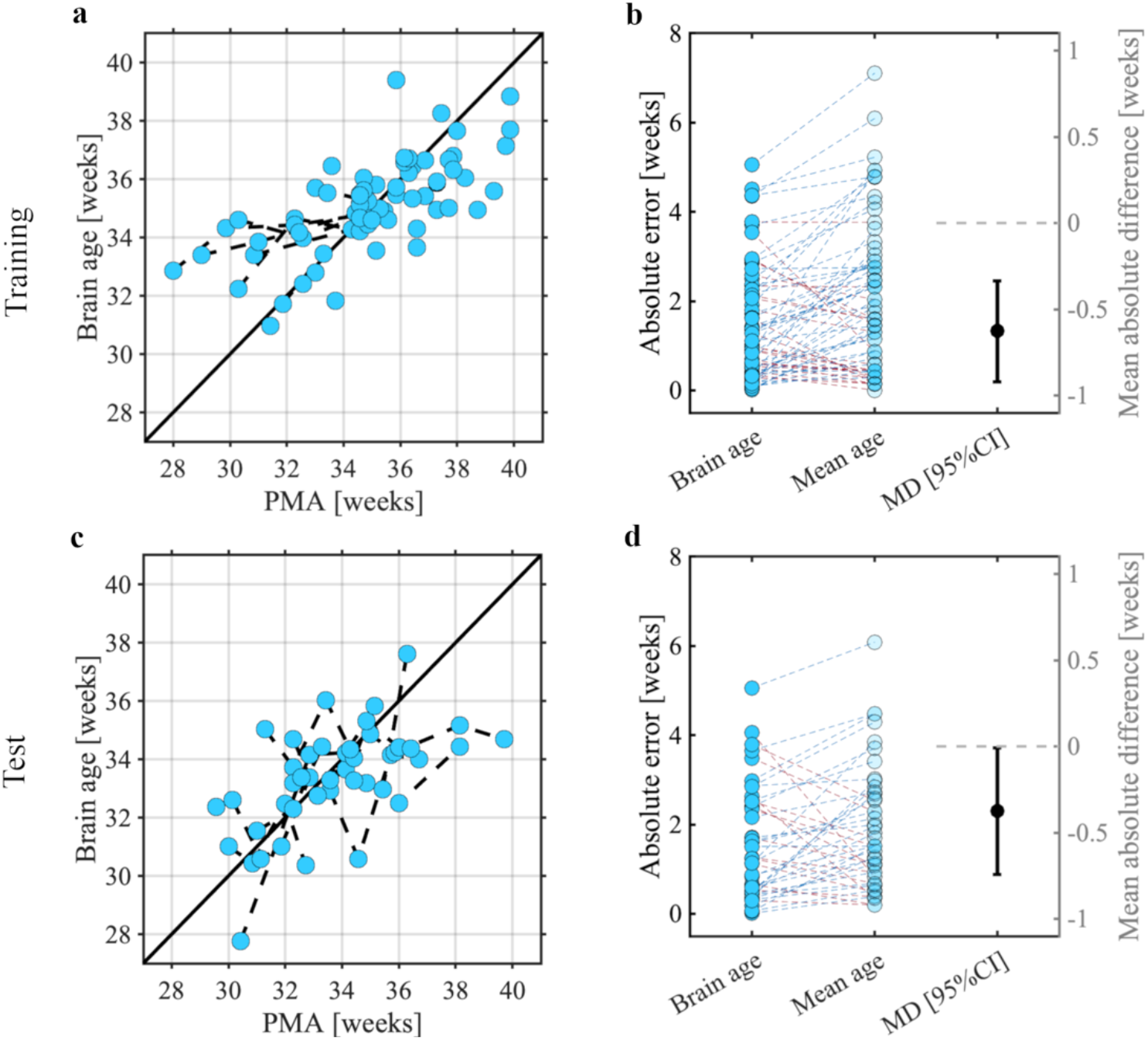
Brain age prediction models and their statistical evaluations for the **a-b)** training and **c-d)** test samples. Panels a and c show the post-menstrual age (PMA) and brain age using leave-one-infant-out cross-validation. Predictions are made from the responses to both visual and tactile stimuli. Each dot indicates a single recording with PMA predicted using the stimulus responses. Dashed lines between dots are infants that took part in multiple recordings. Solid black line indicates perfect prediction. Panels b and d depict the comparison in absolute errors between the Brain age and null model (Mean age) and its mean absolute difference including 95% confidence interval (i.e., MD [95%CI]). Blue dashed lines mean a higher absolute error for the mean age prediction relative to the brain age prediction (i.e., our model performs better than a null model for that recording), and red yield a lower absolute error for the mean age (i.e., our model performs worse than a null model for that recording).

**Figure 4.**
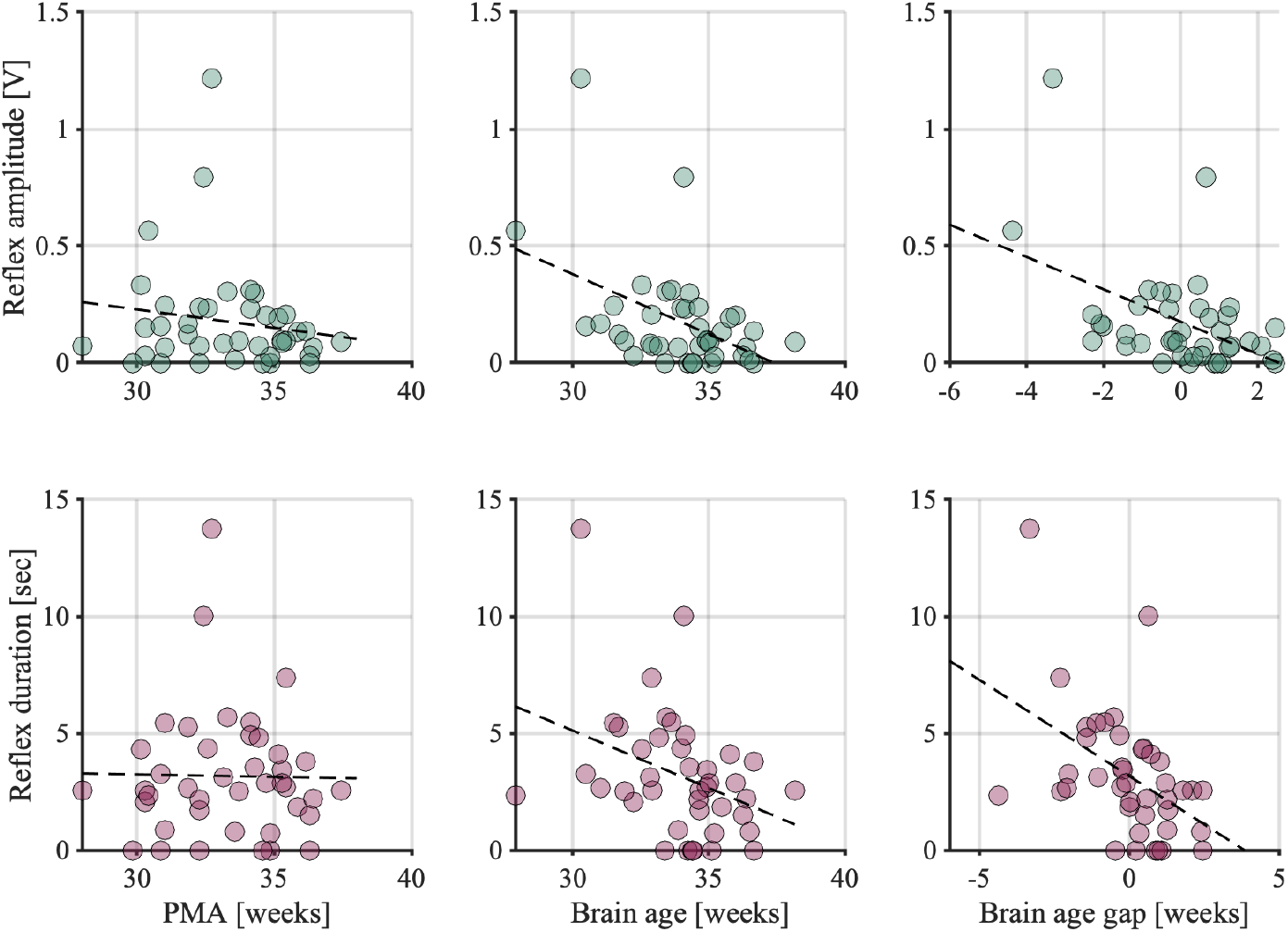
(Brain) age associations with electromyographic reflex responses. The relationship of reflex amplitude and duration following a clinically required heel lance with **a)** post-menstrual age (PMA), **b)** brain age, and **c)** brain age gap. Brain age and brain age gap are derived from the visual+tactile model as presented in Figure 3a. Brain age gap has been adjusted for PMA (see methods). Dashed black graph is the line of best fit.

### Brain age is biologically meaningful – exploratory pilot data 1

For brain age models to be translated into clinical practice, brain age and its difference with PMA, termed the brain age gap, must be biologically and clinically meaningful. Previous studies have shown that the spinally mediated reflex response of an infant to a clinically required painful procedure becomes more refined with age – with shorter duration, smaller amplitude responses (Andrews and Fitzgerald, 1994; Cornelissen et al., 2013; Hartley et al., 2016). This refinement is also well described in animal literature (Fitzgerald et al., 1988; Hathway et al., 2009), and is thought to arise through maturational changes in the sensory nervous system at multiple levels (Brewer and Baccei, 2020). These findings and their theoretical framework suggest that nociceptive reflex withdrawal activity is indicative of maturity of the infant nervous system. Therefore, an infant’s reflex response to noxious stimuli should relate to their brain age if our model is biologically meaningful. In the subset of 32 infants in our study who received a clinically required heel lance at the time of recording, we compared the way they responded to the heel lance with their brain age and brain age gap (i.e., the difference between brain age and PMA). Both brain age and brain age gap are significantly correlated with reflex amplitude (brain age: *r* = -0.45, *p* = 0.002, one-tailed; brain age gap: *r* = -0.46, *p* = 0.001, one-tailed, adjusted for PMA) and duration (brain age: *r* = -0.36, *p* = 0.01, one-tailed; brain age gap: *r* = -0.46, *p* = 0.001, one-tailed, adjusted for PMA).

### Deviations in sensory development may be predictive of later life neurodevelopmental abnormalities – exploratory pilot data 2

Brain age models in infants generally aim to detect atypical development and should ideally be utilised as early indicators of outcomes later in life; hence, linking these two is important. In our sample, infants in the test set are being followed-up at two years of age and assessed using the Bayley Scales of Infant and Toddler Development – Third Edition as part of an ongoing study (see Methods). Five of the infants have already had their two-year follow-up, allowing us to opportunistically investigate the relationship between later life neurodevelopmental outcomes and sensory responses early in life.

Infants were recorded on multiple occasions at approximately one-week intervals. Clear morphological changes with age were observed in response to both visual and tactile stimuli within infants (Figure 5a). Two of the infants had below-average scores for both Language and Motor components (mean score 78 and 76, respectively; both had an average score for Cognitive components: 90), while the other three infants had average or high average scores in all three components (mean across infants of 103, 97, and 100 in Cognitive, Language, and Motor assessments). The two infants with below-average scores had a higher overall MAE of 1.74 weeks, compared to 1.45 weeks for the other three infants. Age predictions in these two infants consistently deviated from their PMA for recordings at older ages (mean gradient of brain age prediction over infants: 0.42 – a gradient of 1 would indicate that brain age is always equal to PMA; Figure 5b), whereas the other three infants showed age predictions that were generally better correlated with PMA (gradient: 0.79; Figure 5b). These deviations at older ages are not the result of noise in the model (Figure S14).

**Figure 5.**
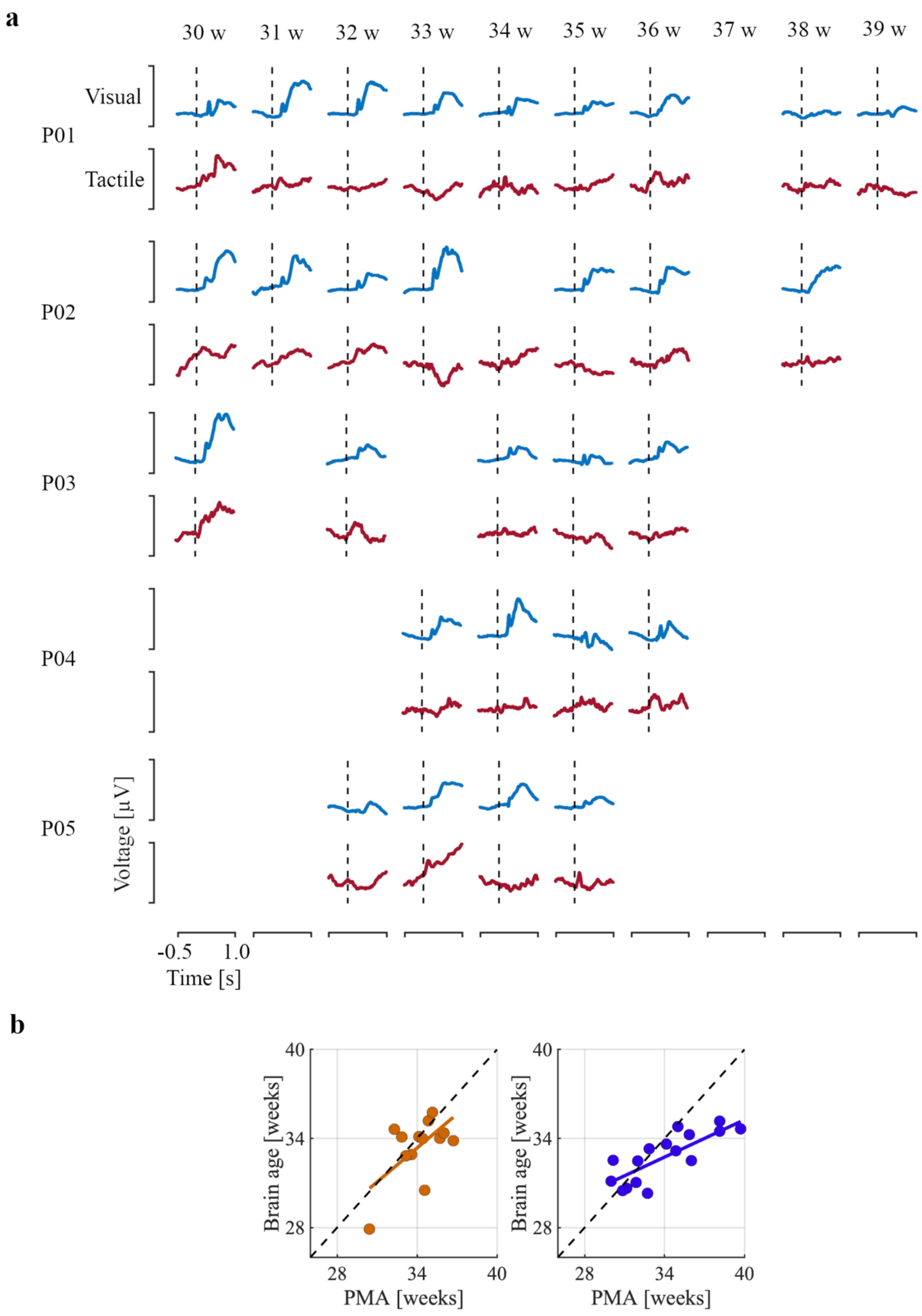
Longitudinal development of the evoked potentials with the brain age predictions of the infants in the test set that have Bayley-III assessments. a) Visual (in blue) and tactile (in red) evoked responses are shown according to post-menstrual age (PMA) at study from 29 to 39 weeks post-menstrual age for each infant indicated by rows P01-P05. Infants were studied approximately once a week during their time in the Newborn Care Unit – infants were born and discharged at variable ages. Vertical dashed line marks stimulus onset. Y-scaling is maximised for each stimulus modality. Infant-specific brain age predictions (on the right) show the predicted PMA using the visual-tactile model for infants with below average (in dark blue, P01 and P02) and average (in dark orange, P03, P04 and P05) neurodevelopmental outcomes. Black lines connect predictions between consecutive sessions. Diagonal dashed black line marks the perfect age prediction. b) Brain age predictions for both groups of infants. Again, orange and blue data are predictions from babies with average and below average Bayley scores. The linear regression is the mean of the line of best fit over infants.

## Discussion

We aimed to quantify standardised multisensory brain responses in infants aged 28 to 40 weeks PMA and exploit these standardised responses to predict the brain age of the infants. We applied our neurodynamic response function (NRF) analysis approach to identify distinct stimulus-evoked brain responses to visual and tactile stimuli. This data-driven approach revealed four stimulus-specific NRFs in response to the visual stimulus and two in response to the tactile stimulus. Brain age could be accurately predicted from the magnitudes of these NRFs, and we validated this model in an independent test set. Brain age (gap) was correlated with the magnitude and duration of the reflex withdrawal response to a heel lance, suggesting that deviations in brain age are biologically and clinically meaningful. Moreover, in a subset of the test set with neurodevelopmental outcome at two years of age, we show that sensory evoked brain age deviated from PMA in infants with below average outcome in the Bayley Scales of Infant and Toddler Development at two years of age, suggesting that sensory-evoked potentials (and our brain age model) are predictive of later life outcome.

The brain age model comprising sensory-evoked responses captures the rapid structural and functional development of the neurosensory system of (premature) infants. The neural architecture to process sensory stimuli at the cortex is established from around the start of the third trimester (Colonnese and Khazipov, 2012), with thalamocortical connections initially via the transient subplate (Kostović et al., 2014; Kostović and Judaš, 2010). Brain activity in this period is characterised by intermittent bursts of activity including delta brush activity (higher frequency neural oscillations nested within a delta wave) (Khazipov et al., 2004), which can occur spontaneously or be evoked by stimuli (Milh et al., 2007; Whitehead et al., 2017). As PMA increases, delta brush activity begins to disappear and evoked brain activity with high frequency waveforms emerge (André et al., 2010; Niemarkt et al., 2011). The disappearance of delta brush activity is apparent in sensory-evoked activity (Chipaux et al., 2013; Colonnese and Khazipov, 2012; Fabrizi et al., 2011; Hartley et al., 2016; Kato and Watanabe, 2006; Mercuri et al., 1994; van der Vaart et al., 2022) but the timepoint at which the transition from delta brush to modality-specific evoked potentials occurs may be dependent on stimulus modality (Colonnese and Khazipov, 2012). Consistent with these previous studies, we identified age-dependent changes in the stimulus-evoked responses. In our study, delta waves are particularly captured by the visual NRFs 1 and 2 and tactile NRF 1. For all three NRFs, these responses occurred mostly in younger babies as expected. Higher frequency waveforms were apparent in the second NRF in response to tactile stimulation and in NRFs 3 and 4 in response to visual stimulation.

A wide range of brain age models has been developed to trace the brain development of premature infants, encompassing structural connectivity (Brown et al., 2017; Kawahara et al., 2017), morphological (Liu et al., 2021) and electrophysiological data. For the latter, brain age models have previously been constructed in preterm infants using resting state EEG recorded brain activity (Ansari et al., 2023; Lavanga et al., 2018; O’Toole et al., 2016; Pillay et al., 2020; Stevenson et al., 2017). Although the MAE achieved by our model is not as accurate as some resting state models (e.g., MAEs of approximately 1 week were achieved by Ansari et al. (2023) and Liu et al. (2021)), compared to these existing brain age models, our model has the advantage that it was constructed using electrophysiological responses of approximately 10 visual and 10 tactile stimuli from every recording. We applied stimuli with an inter-stimulus interval of approximately 10 seconds; however, it may be possible to present them at shorter latencies. Nevertheless, this means that brain age predictions can be made based on approximately 5 minutes of recording. Current brain age models utilising ongoing resting state activity require at least 20 minutes of EEG data (Ansari et al., 2023). Implementing sensory evoked responses into brain age models has the potential to lower the requirements on the amount of data that needs to be acquired in a busy clinical environment. Integrating such data will also provide information about the integrity of sensory pathways and will add value to existing resting state models. Indeed, it may be possible to combine the sensory and resting state brain age models, which potentially allows for a more comprehensive understanding of both the underlying functional brain architecture and sensory responses to environmental stimuli.

Our model used responses to both visual and tactile stimuli, which performed better than either stimulus individually. This could in part be due to the smaller numbers of features used in the single-stimulus models compared with the multi-modal model. Further work could explore the use of other features such as the latency to the response, which is known to be age dependent (Schwindt et al., 2018; Taylor et al., 1987). Nevertheless, it makes intuitive sense that including multimodal responses will improve accuracy and future work should also consider including responses to other stimuli such as auditory and noxious.

For brain age models to be useful, the infant’s brain age (or the deviation between their PMA and brain age) should be biologically meaningful rather than just noise generated by the model (i.e., errors made in the prediction due to non-biological sources such as differences in head size). Thus, brain age should be correlated with variables indicating the integrity of the neurosensory system. To test this in an example situation, we compared the infant’s brain age with the magnitude of the spinally mediated reflex withdrawal to a noxious stimulus. We chose reflex withdrawal measured with electromyography rather than the EEG-recorded noxious evoked brain activity to the stimulus as the EEG response may be well-correlated with visual and tactile-evoked derived brain age due to EEG intrinsic noise factors such as electrode placement rather than biologically meaningful factors. In young rat pups, reflex withdrawal to noxious stimuli is uncoordinated and exaggerated compared with adult animals (FitzgeraId et al., 1988; Hathway et al., 2009; Holmberg and Schouenborg, 1996). The change in reflex withdrawal over the first few weeks of postnatal life corresponds to the development of descending inhibition, a reduction in cutaneous receptive fields, and changes in innervation and activity of the spinal cord dorsal horn (Brewer and Baccei, 2020; Fitzgerald, 1985; Holmberg and Schouenborg, 1996; Koch and Fitzgerald, 2013). In line with the animal literature, in preterm infants’ reflex withdrawal decreases in magnitude and duration with age, and the threshold for the response increases (Andrews and Fitzgerald, 1994; Cornelissen et al., 2013; Fitzgerald et al., 1988; Hartley et al., 2016). Here, we found that an infant’s brain age and brain age gap are correlated with the magnitude of the reflex withdrawal. From the strong basis of animal literature, it is expected that the reflex withdrawal is related to the maturity of the nervous system. Thus, this gives support to suggest that brain age is biologically meaningful. Moreover, it may be clinically useful in this scenario as brain age may lead to a better understanding of infants’ responses to painful procedures and so could be, for example, useful for testing analgesics. Further research in this area is warranted (Moultrie et al., 2017; Slater et al., 2020).

In the test set, we found initial support that our sensory brain age predictions are associated with neurodevelopmental outcomes at two years of age, whereby a higher brain age gap (i.e., the difference between PMA and predicted age) was correlated with the poorer neurodevelopmental outcome as defined using Bayley-III scores. This is in line with previous results from resting state models (Ansari et al., 2023; Pillay et al., 2020; Stevenson et al., 2020). Future studies should examine how the brain age gaps from abnormal sensory-evoked responses relate to the neurodevelopmental outcomes in larger samples. The longitudinal recordings included here provide evidence that an infant’s brain age may begin to deviate from PMA at certain time points which are likely individualistic. Longitudinal follow-up provides an opportunity to investigate factors that lead to altered neurodevelopment and identify possibilities for intervention.

To conclude, we present a brain age model constructed using sensory-evoked responses in premature infants. This brain age model accurately predicts age, including in an independent test set, and sensory-evoked brain age deviated from PMA in infants with below-average neurodevelopmental outcome. Moreover, brain age (gap) is correlated with spinally mediated reflex withdrawal responses, suggesting it is biologically meaningful. Compared with current models constructed using resting state EEG, it requires only a limited number of sensory evoked potentials (on average 20 epochs of 1 second duration), which could be regularly assessed at the cot-side. Recording these EEG responses can be achieved with 5 minutes of data collection. Assessment of neurological function and the integrity of sensory pathways in premature infants is essential for prognostication of later life outcome and the provision of early targeted interventions.

## Material and methods

### Participants and study design

All infants were selected from a research database, containing the data acquired during other experimental protocols, including those presented in previous reports (Green et al., 2019; Hartley et al., 2017; Schmidt Mellado et al., 2022). These data were collected between 2012 and 2023 at the John Radcliffe Hospital, Oxford University Hospitals NHS Foundation Trust, Oxford, United Kingdom. Studies were approved by the National Research Ethics Service (ethics references: 12/SC/0447; 19/LO/1085; 11/LO/0350). Parents or legal guardians provided verbal and written consent before participation in the research studies. All study protocols complied with the Declaration of Helsinki and guidelines on Good Clinical Practice.

Infants were included in the analysis if they had brain activity responses recorded following either visual or tactile stimuli. An exclusion criterion was intraventricular haemorrhage (IVH) grade 3 or 4. Infants were divided into a training and test sample. A total of 101 recordings were identified from the database and were labelled as the training sample. Seventy-nine of these recordings included visual stimuli and 95 recordings included tactile stimuli. These were recordings from 82 unique infants – 70 infants were recorded on one test occasion only, 6 infants were recorded twice, 5 infants were recorded on three test occasions, and 1 infant was recorded on four separate occasions. Infants were born between 23- and 40-weeks’ gestation and were aged between 28- and 40-weeks PMA at the time of the test occasion. Infants in the independent test sample were all recruited as part of the ongoing ‘Breathing and Brain Development’ study (https://www.hra.nhs.uk/planning-and-improving-research/application-summaries/research-summaries/breathing-and-brain-development-version-10/). All infants recruited as part of this study up to February 2023 were included in the test sample, giving a total of 14 infants recorded on 65 occasions. Both visual and tactile stimuli were applied in 57 recordings. PMA in the test sample ranged between 29 and 40-weeks’ gestation. Full demographic details are provided in Table 1.

**Table 1.** Reported values are mean (range) or number (%) of babies or recordings. All demographic details apart from post menstrual and postnatal age (PMA and PNA, respectively) are provided per infant (PMA and PNA are computed for every recording).

| **Factors** | *Training sample* | *Test sample* |
| --- | --- | --- |
| *Age* |  |  |
| PMA at recording (weeks) | 34.8 (28.0-39.9) | 33.7 (29.6-39.7) |
| Gestational age (weeks) | 32.7 (23.6-39.7) | 29.7 (28.1-32.6) |
| PNA at recording (weeks) | 2.9 (0.0-11.4) | 4.2 (0.6-10.7) |
| Birthweight (g) | 1,929 (630-4,525) | 1,272 (635-2,120) |
| *Sex* |  |  |
| Females | 37 (45.1) | 4 (28.6) |
| Males | 45 (54.9) | 10 (71.4) |
| *Mode of delivery* |  |  |
| Normal vaginal delivery | 24 (29.3) | 2 (14.3) |
| Vaginal breech | 3 (3.6) | 0 (0.0) |
| Vaginal assisted (ventouse/forceps/kiwi) | 5 (6.1) | 2 (14.3) |
| Elective C-section | 10 (12.2) | 1 (7.1) |
| Emergency C-section/C-section in labour | 40 (48.8) | 9 (64.3) |
| *Apgar scores* |  |  |
| Apgar at 1 minute | 7.6 (1-10) | 6.4 (1-10) |
| Apgar at 5 minutes | 9.2 (3-10) | 8.8 (5-10) |
| Apgar at 10 minutes | 9.8 (6-10) | 9.6 (8-10) |

## Data acquisition

### EEG recordings and stimuli

SynAmps RT 64-channel headbox and amplifiers (Compumedics Neuroscan, Compumedics Limited, Victoria, Australia) and CURRYscan7 neuroimaging suite (Compumedics Neuroscan, Limited, Victoria, Australia) were used to record the EEG data at a sampling rate of 2 kHz (EEG data during two recordings of the training sample were acquired at 1 kHz and resampled to 2 kHz). The EEG channel configuration included channels Cz, CPz, C3, C4, FCz, Oz, T3, and T4. Channel Fz was selected as the reference electrode while FPz served as the ground electrode. The electrode array configuration was in line with the international 10-20 system. To optimise contact with the scalp, the skin was gently rubbed with EEG preparation gel (NuPrep gel, D.O. Weaver and Co., Aurora, USA) prior to electrode placement. EEG conductive paste (Elefix EEG paste, Nihon Kohden, Tokyo, Japan) was used with disposable Ag/AgCl cup electrodes (Neuroline, Ambu, Ballerup, Denmark).

A series of visual and tactile stimuli were presented to the infants in a pseudo randomised order, with the researcher deciding which stimuli to present first. The visual stimulus consisted of a light flash presented using a Grass LED light (Maxima-84 Hybrid, Manfotto, Italy) or Lifelines Photic Stimulator (Lifelines Ltd.; flashing frequency: 10 Hz; intensity level: 4, which approximates 514 lm). The former stimulus type was presented at 50 cm from the infant’s eyes (8 recordings); the latter at a distance between 15 and 30 cm (71 recordings in the training sample and 57 recordings in the test sample. The light was positioned at less than 30 cm if there was limited space in the incubator). All visual stimulation types were automatically annotated on the EEG at the time of the recording. Infants received a median number of 12 (interquartile range (IQR) = 13) visual stimuli in the training sample (*n* = 79) and 10 (IQR = 1) in the test sample (*n* = 57), with median interstimulus intervals of 11.0 s (IQR = 1.8 s) and 11.6 s (IQR = 3.2 s) per recording, respectively.

For the tactile stimulus, a researcher gently touched the heel of the infant using a modified tendon hammer. This tendon hammer recorded the applied force via a built-in transducer (Brüel & Kjær, Type 8001, Denmark) used to time-lock the stimulus with the EEG recording (Worley et al., 2012). Infants received a median number of 12 (IQR = 15) and 10 (IQR = 1) tactile stimuli in training (*n* = 95) and test (*n* = 57) sample, respectively with interstimulus intervals of 11.0 s (IQR = 2.7 s) and 11.9 s (IQR = 4.0 s) per recording.

A researcher made real-time resting state activity annotations during recordings when no stimuli were applied and the infant was quietly awake or asleep, and not moving. The resting state activity served as reference condition in the statistical contrasts of the cluster-based permutation and NRF magnitude comparisons (see below). A median of 16 (IQR = 11) and 11 (IQR = 10) resting state annotations were made per recording in the training and test sample, respectively.

### Electromyographic recordings and clinically required heel lance

Bipolar electromyographic (EMG) electrodes (Ambu Neuroline 700 solid gel surface electrodes) were attached to the biceps femoris of the infant’s leg ipsilateral to the site of stimulation and recorded using the same recording system as for the EEG electrodes. Heel lances were performed in infants if they clinically required a blood test at the time of the test occasion. The heel lance was time-locked to the EMG (and EEG) recordings using an event detection interface and accelerometer (Worley et al., 2012).

### Neurological outcomes

To assess how the brain age model outcomes relate to behavioural and neurological developmental outcomes at 24-months follow-up age, Bayley Scales of Infant and Toddler Development – Third Edition (Bayley-III) were obtained from five (out of the 14) infants of the test set at the time of this report (note that the other nine infants will have Bayley assessments once they reach two years of age as part of an ongoing study). Bayley assessments were taken at a mean age of 2 years, 3 months, and 26 days (minimum: 2 years, 2 months, 25 days, and maximum: 2 years, 4 months, 22 days). We report composite scores for Motor, Cognitive and Language outcomes.

## Data analysis

### EEG pre-processing

We focused analysis on channels Cz for tactile-evoked activity and Oz for visual-evoked responses, in line with analysis from a previous study that comprised parts of the dataset used in the current study (Schmidt Mellado et al., 2022). From a neuroanatomical point of view, channels Cz and Oz overlay the primary somatosensory and visual cortices, respectively, and maximal amplitude responses are expected at these electrodes.

EEG data were processed using custom-made scripts in MATLAB (ver. 2022b; MathWorks Inc., Natick, USA) together with Brainstorm (ver. 3) (Tadel et al., 2011) and EEGLAB (ver. 2022.1) (Delorme and Makeig, 2004). Continuous EEG data were filtered with low-pass (Hamming windowed-sinc FIR filter with pass-band edge at 30 Hz and cut-off frequency at 33.75 Hz) and high-pass filters (Hamming windowed-sinc FIR filter with pass band edge at 0.1 Hz and cut-off frequency at 0.05 Hz). To derive NRFs, the EEG was further filtered with a low-pass filter of 12 Hz (cut-off frequency at 13.5 Hz) because instantaneous amplitude representations showed that spectral power drops above 12 Hz (Figure S15). This is expected, as EEG activity is dominated by lower frequency (i.e., delta, theta and alpha) activity in premature infants (André et al., 2010), and filtering the activity enables clear characterisation of the waveforms within the evoked response. However, when examining how the magnitudes of these NRFs change with age and for the brain age prediction models, the EEG data were filtered between 0.1 and 30 Hz.

EEG was epoched from 1 s before until 1 s after stimulus onset and visually inspected around the stimulus. Individual epochs (of all channels) were rejected when the amplitude in the pre-stimulus window exceeded +/-150 µV between –1 and 0 s (i.e., unstable baseline). This led us to reject an average of 3 ± 5, 1 ± 2, and 2 ± 3 epoch(s) (median ± interquartile range) per recording in the training sample for the resting state, tactile, and visual annotations, respectively. In the training sample, we retained an average of 11 ± 11, 12 ± 11, and 11 ± 10 epochs for further analysis in the corresponding test conditions highlighted above. In the test sample, 0 ± 3, 0 ± 1, and 1 ± 2 epochs were excluded, with a final 10 ± 9 resting state, 10 ± 1 tactile, and 10 ± 1 visual epochs included for the analysis.

We also excluded recordings where fewer than five responses of a certain stimulus type were available. This led us to reject 12 stimulus conditions (4 visual and 2 tactile in the training sample and 2 visual and 4 tactile in the test sample). This meant that one baby was excluded in the training sample as both visual and tactile stimuli were rejected. For the resting state activity, if there were only five or fewer events available, we created ten new events by adding them with a time interval of 10 s prior to the first annotated resting state event. Resting state events were added for 15 recordings (7 in the training sample and 8 in the test sample). For one baby with a single recording, all resting state annotations were removed because of excessive amplitudes and was thus excluded from further analysis (this infant also had fewer than 5 tactile stimuli). Overall, for the training sample we included 74 recordings with visual-evoked responses and 93 recordings with tactile-evoked responses (these were 98 unique recordings from 80 babies). For the test sample, a total of 65 recordings comprising 55 visual and 53 tactile potentials were included from 14 babies.

### Developing response functions for visual- and tactile-evoked brain activity

NRFs were derived from the training sample. We first computed recording-specific response averages and then age-weighted averages of temporal alignment; next, we identified time periods with significant stimulus-evoked activity using cluster-based permutation testing; and finally, we identified waveforms characteristic of the stimulus response using principal component analysis (PCA).

EEG data in the time window of 0 to 1 s post-stimulus were baseline corrected to the time window of -0.5 to 0 s pre-stimulus. Demeaned responses were pooled over epochs for each stimulus modality to create recording-specific averages. These EEG responses were temporally shifted to an age-weighted response using Woody filtering to adjust inter-recording differences in response latency (Woody, 1967). The age-weighted responses were constructed by assigning a weight of 1 or lower to each recording depending on PMA. A Gaussian window with a full width at half maximum of 27 days determined the weights of neighbouring PMAs. Age differences of more than 28 days with the recording of interest received a weight of 0. Age-weighted responses were computed by scaling every recording-specific average with its weight and taking the sum of these responses divided by the sum of the weights. We then Woody filtered each recording-specific average to an age-weighted response (maximal jitter: 0.05 s). The time-shifted responses for each recording were used in the cluster-based permutation testing and PCA and so both resting state and stimulus responses were Woody filtered within the age-dependent responses (enabling fair comparisons between the stimulus response and resting state).

To evaluate in which time windows the stimulus amplitude significantly differed from the resting state amplitude in the time window of 0 to 1 s post-stimulus, we applied cluster based permutation testing (Maris and Oostenveld, 2007). This nonparametric approach iteratively performs sample-wise paired t-tests between stimulus and resting state responses for every session. Time samples exceeding a pre-defined *t*-value threshold (here, set to 97.5 percentile of the *t*-distribution and degrees of freedom minus 1) were defined as significant activity, with adjacent significant samples defined as a cluster. Clusters were defined as significant (α-level of 0.05) by comparing with the distribution of clusters obtained from 1,000 permutations of the data (stimulus and resting state traces were permuted in a paired way and partitioned into one of the two conditions). To compute the NRFs, data were trimmed to the time windows of 0 to 0.3 s and 0.1 to 1 s for the tactile and visual responses respectively (Figure S2), which was around the significant clusters and allowed for characterisation of the full waveform associated with the significant cluster. These trimmed responses were used as input for the PCA.

To derive the NRFs, PCA decomposed the (1) visual and resting state, and (2) tactile and resting state responses into sets of covarying waveforms that explained most variance across the evoked brain responses. We normalised the recording-specific averages to unit vectors, meaning that the PCs reflected morphological changes of the stimulus response across age rather than being dominated by inter-recording amplitude differences. Responses were normalised to their Euclidian norm over the entire time window. We extracted the number of PCs that could explain more than 95% of the variance, which yielded seven and four PCs for the visual and tactile responses. To identify which of these components were indicative of stimulus response we compared the weights of the components between resting state and stimulus responses by fitting each temporal PC-component to the non-normalised stimulus and resting state activity using linear regression (Table S1). Mean stimulus and resting state magnitudes were statistically compared using two-sided paired t-tests (*p* < 0.05). The temporal components of the significant PCs were taken as the NRFs.

### Characterising developmental changes in NRFs

The broad-band filtered (0.1-30 Hz) stimulus-evoked EEG responses were Woody-filtered to the NRFs in the time window of 0-1 s post-stimulus (jitter: 0.05 s). This minimised the latency differences between recording-specific responses and NRFs. The magnitude of the NRF for each recording was determined by linearly regressing the NRFs to the Woody-filtered responses and calculating the slope coefficients (akin to the process used in fMRI when calculating the beta coefficient at each voxel compared with the haemodynamic response function). For each NRF, relationships between PMA with the training- and test-sample magnitudes were quantified by fitting generalised linear regressions with identity link functions to the averaged NRF magnitudes for every week over PMA. *p*-values were used as a guide and no correction was made for multiple comparisons.

### Predicting brain age using support vector regression

To predict the brain age of infants during each recording, we used support vector regression (SVR) with a linear kernel function. Errors and allowed margin from these errors were set to 0.15 and 1, respectively, which are parameters defined based on the interquartile range of the PMA. The L1 soft-margin minimisation was used as solver. The model was implemented in MATLAB using the fitrsvm function (version 2022b; MathWorks).

Predictor variables were the NRF magnitudes for each recording with all NRFs. In the training sample, we created three models with different predictor variables. The first model contained the visual NRFs defined on Oz and tactile NRFs on Cz. The response variable was PMA for the 98 recordings of the 80 unique infants who had responses to either visual or tactile stimuli. A model was also trained with only visual responses and separately with only tactile responses (see supplementary material).

We used leave-one-infant-out cross-validation to assess the model performances in the training set, calculating the mean absolute error (MAE) between the PMA and brain age, with 95% confidence intervals estimated from 10,000 bootstrap samples. Significance was obtained using one-tailed testing using permutation testing as provided in FSL’s PALM (Winkler et al., 2014). Permutations were limited to pre-defined exchangeability blocks because of the multiple recordings for every infant (Winkler et al., 2015). Lastly, in addition to the reported MAE, the true model output was compared to a model which predicts the mean PMA over recordings (with the mean age calculated using leave-one-infant-out). We report mean absolute differences, confidence intervals, and *p*-values between the true models and mean PMA models.

Finally, we applied the training sample model (calculated with all training data) to predict the PMA of babies in the test sample using their NRF magnitudes as input. Model performance was assessed by estimating the MAE and its associated 95% confidence intervals. Model significance was estimated by comparing the actual model to a null model predicting the mean age of the test sample. We note that we first derived the NRFs and brain age model in the training set before studying the test set, results here are shown together for ease of comparison.

### Brain age model application to electromyographic reflexes to noxious stimuli

To demonstrate that the brain age model we designed is neurobiologically meaningful, we examined how the predicted brain ages correlated with the EMG-recorded withdrawal reflexes in response to painful procedures. In both the training and the test set, there were 40 recordings which we predicted brain age with, including the visual+tactile model and EMG recorded at the biceps femoris during a clinically required heel lance. EMG recordings were filtered from 10 to 500 Hz (Hamming windowed-sinc FIR filter with cut-off frequencies at 8.75 and 562.5 Hz), with a notch filter at *k*\*50 Hz (with *k* = 1, 2, …, 10), epoched from 5 seconds before the stimulus until 15 seconds afterwards, and rectified. From the rectified EMG, we defined reflex duration and amplitude using the methods described in Hartley et al. (2016), which uses an automated algorithm to detect the start and end of the reflex. Epochs were visually inspected and rejected if there was movement in the baseline period precluding the identification of the start of the reflex. A total of 8 recordings were rejected, leaving 32 recordings in the analysis. Reflex amplitude and duration were then linearly correlated with PMA, brain age, and brain gap (i.e., the difference between brain age and PMA). The brain age gap was adjusted for PMA by creating a linear model that predicts the brain age gap from the PMA. The residuals of this linear model were taken as the adjusted brain age gap values.

## Data and code availability

The data that support the study findings are available from the corresponding author upon reasonable request. Due to ethical restrictions, it is appropriate to monitor access and usage of the data since it includes highly sensitive information. Data sharing requests should be directed to. The NRFs and codes underpinning the brain age model are available on GitLab: https://gitlab.com/paediatric_neuroimaging/sensory-brain-age-model.

## CRediT authorship contribution statement

**Coen S. Zandvoort:** Methodology, Software, Formal analysis, Writing – Original Draft, Writing – Review & Editing, Visualisation; **Marianne van der Vaart:** Methodology, Data Curation, Writing – Review & Editing; **Shellie Robinson:** Investigation, Writing – Review & Editing; **Fatima Usman:** Investigation, Writing – Review & Editing; **Gabriela Schmidt Mellado:** Investigation, Data Curation, Writing – Review & Editing; **Ria Evans Fry:** Investigation, Writing – Review & Editing; **Alan Worley:** Methodology, Writing – Review & Editing; **Eleri Adams:** Supervision, Writing – Review & Editing; **Rebeccah Slater:** Supervision, Writing – Review & Editing; **Luke Baxter:** Methodology, Data Curation, Writing – Review & Editing; **Maarten de Vos:** Methodology, Writing – Review & Editing; **Caroline Hartley:** Conceptualisation, Funding acquisition, Methodology, Data Curation, Writing – Original Draft, Writing – Review & Editing, Supervision

## Acknowledgements

The authors would like to thank all parents and infants involved in the studies and staff at the John Radcliffe Hospital in Oxford. This study was funded by a Wellcome Trust/Royal Society Sir Henry Dale Fellowship (grant number: 213486/Z/18/Z), awarded to CH. MvdV, LB and RS are funded a Wellcome Trust Senior Research Fellowship awarded to RS (grant number: 207457/Z/17/Z). FU is funded by the Commonwealth Scholarship Commission. This research was funded in whole, or in part, by the Wellcome Trust 213486/Z/18/Z. For the purpose of Open Access, the author has applied a CC BY public copyright licence to any Author Accepted Manuscript version arising from this submission.

## Supplementary material Supplementary text Results

### Visual and tactile models

Besides the brain age model that was constructed from both visual and tactile magnitudes, we also made single-modality models using either visual or tactile magnitudes as input. For the training set, mean absolute errors (MAE) were higher compared to the visual-tactile model but age prediction performance was still significantly different from the average age model (visual only: MAE: 1.77 weeks with 95% at [1.46; 2.16], *p* = 0.0003; Figure S12, tactile only: MAE: 1.67 weeks with 95% at [1.35; 2.03], *p* = 0.0024; Figure S13). However, in the independent test sample, the single-stimulus models of visual and tactile responses, age prediction was not significantly different from the average age model (visual - MAEs = 1.75 weeks with 95% at [1.51, 2.03], *p* = 0.0001; Figure S12; tactile MAE = 1.77 weeks with 95% at [1.44; 2.14], *p* = 0.0005; Figure S13).

### Deviations in sensory development may be predictive of later life neurodevelopmental abnormalities – exploratory pilot data 2

To exclude that the results as presented in Figure 5 were related to bias in our model (particularly at older ages, where the error in the training set is greater than for infants at approximately 34 weeks and the brain age may be underpredicted, Figure 3a), we recalculated the gradients without recordings from when the infants were older than 37 weeks. Restricting the age range, the mean gradients for the two infants with below average Bayley’s outcomes equalled 0.48 and for the three infants with average Bayley’s outcomes equalled 0.79. Secondly, we estimated the bias in the training set and removed this from the test set (Figure S14). After bias removal, whilst the gradients of the brain age trajectories were closer to 1 (gradient of 1.00 for infants with below average Bayley’s and 1.37 for those with average outcomes, Figure S14) but the MAE was still higher in the infants with below average outcome (1.91 compared with 1.57 weeks), supporting the finding that brain age may be indicative of later life outcome.

## Supplementary tables

**Table S1.** NRF magnitudes and statistical comparisons between the stimulus responses and resting state activity. Statistics belong to the paired comparisons between stimulus and resting state magnitudes. Explained variance column contains the percentage of variance that the NRF can explain of the event-related data. NRF: neurodynamic response function; std: standard deviation.

|  | Neurodynamic response function | Explained variance (%) | Stimulus magnitudes (mean±std) | Resting state magnitudes (mean±std) | t-value | Degrees of freedom | p-value |
| --- | --- | --- | --- | --- | --- | --- | --- |
| Training set | Visual NRF1 | 56.5 | 157.6 ± 161.5 | 11.7 ± 60.1 | 7.06 | 73 | <0.0001 |
|  | Visual NRF2 | 24.7 | 90.0 ± 123.5 | 7.9 ± 48.1 | 5.04 | 73 | <0.0001 |
|  | Visual NRF3 | 2.4 | 67.2 ± 179.1 | -5.6 ± 37.1 | 3.59 | 73 | 0.0006 |
|  | Visual NRF4 | 1.4 | 52.1 ± 164.9 | -8.0 ± 35.4 | 3.15 | 73 | 0.002 |
|  | Tactile NRF1 | 46.7 | 12.0 ± 37.4 | 0.5 ± 12.3 | 3.35 | 92 | 0.001 |
|  | Tactile NRF2 | 36.4 | 20.7 ± 27.2 | 1.2 ± 6.5 | 6.53 | 92 | <0.0001 |
| Test set | Visual NRF1 | N/A | 304.2 ± 266.0 | -0.3 ± 70.1 | 8.14 | 54 | <0.0001 |
|  | Visual NRF2 | N/A | 250.9 ± 185.3 | 13.7 ± 69.6 | 8.29 | 54 | <0.0001 |
|  | Visual NRF3 | N/A | 54.6 ± 155.0 | 4.2 ± 49.6 | 2.40 | 54 | 0.02 |
|  | Visual NRF4 | N/A | 240.2 ± 297.6 | -6.0 ± 39.2 | 5.89 | 54 | <0.0001 |
|  | Tactile NRF1 | N/A | 12.6 ± 25.7 | -0.3 ± 14.1 | 3.53 | 52 | 0.0009 |
|  | Tactile NRF2 | N/A | 12.3 ± 13.9 | 1.1 ± 6.6 | 5.02 | 52 | <0.0001 |

**Table S2.**
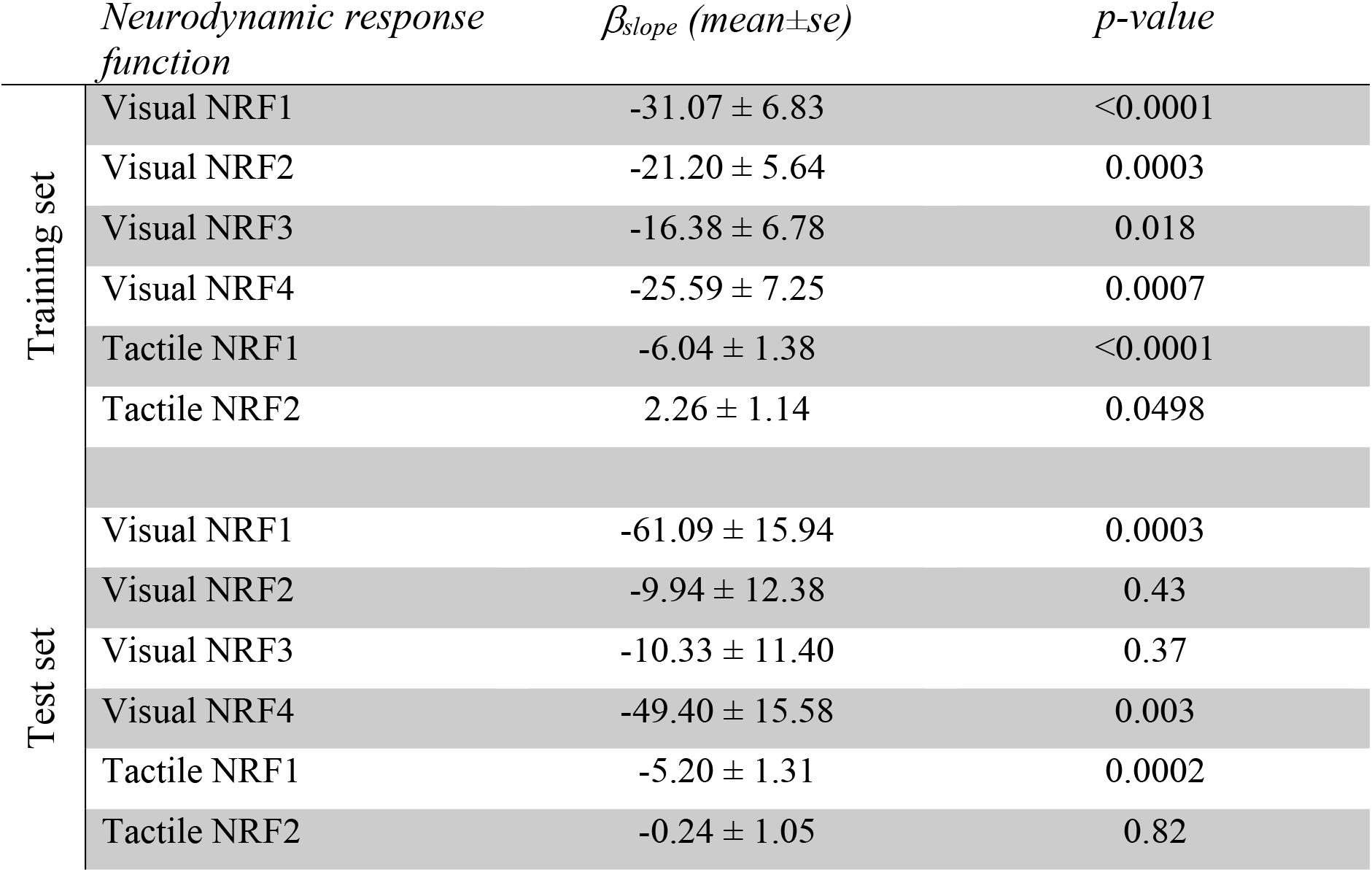
Linear regression slope coefficients for each of the neurodynamic response functions (NRFs) of the training and test set. Regression models predicted stimulus magnitudes from the post-menstrual age (PMA). se: standard errors.

## Supplementary figures

**Figure S1.**
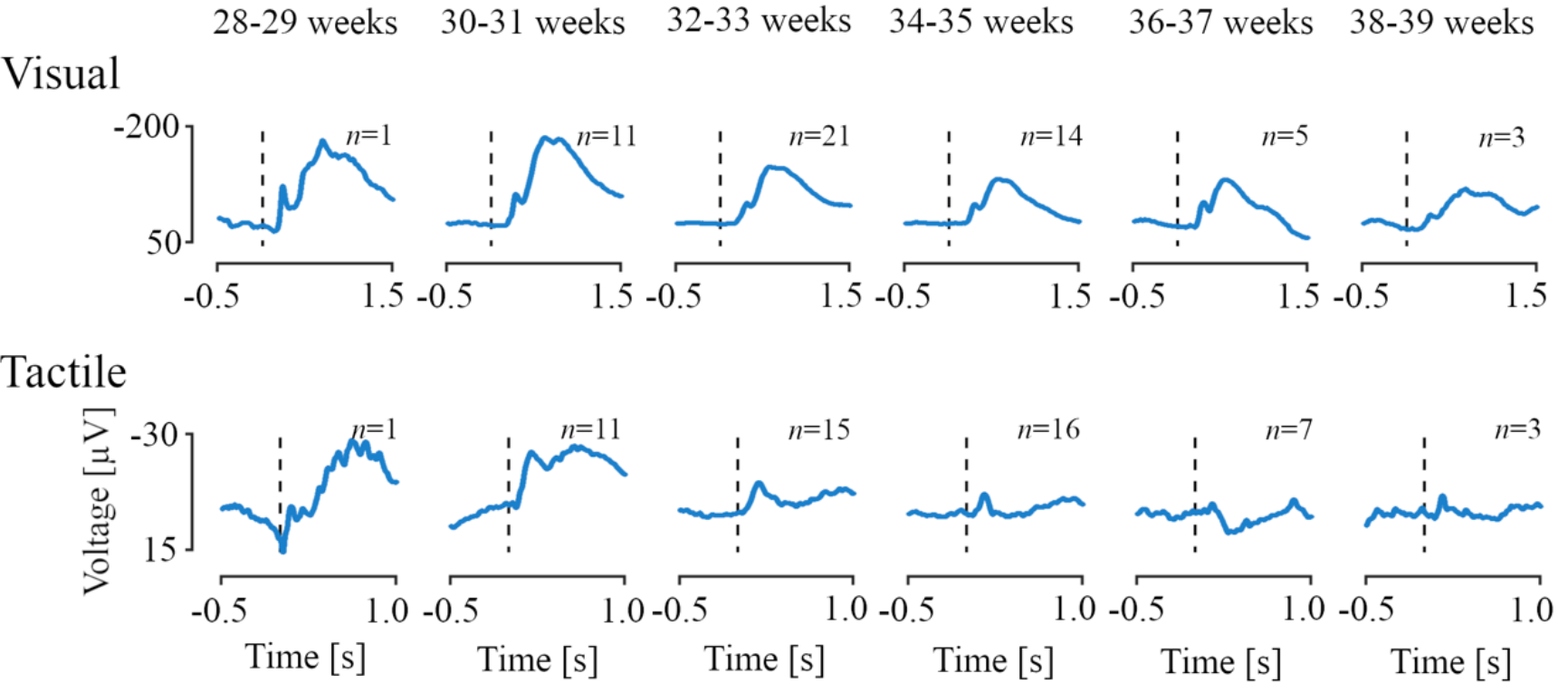
Stimulus-evoked electroencephalographic potentials according to infant age. Age-dependent evoked potentials for two-weeks intervals between 28 to 40 weeks of post-menstrual age for the visual and tactile stimuli at channels Oz and Cz, respectively, for the test set (see Figure 1 for the training set). Woody filtering aligned the responses to their age-weighted averages. Vertical dashed lines correspond to the stimulus onset.

**Figure S2.**
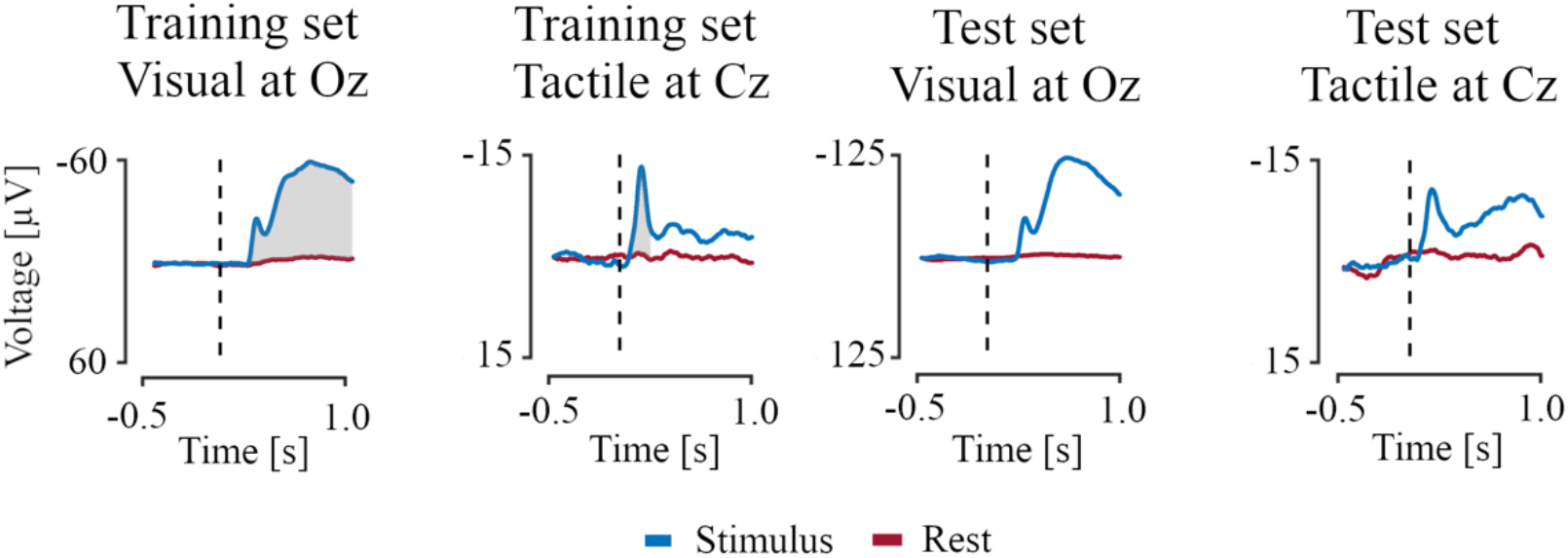
Grand average means of the stimulus response (in blue) and resting state (in red). Grey areas depict the time windows where the stimulus and resting state means are significantly different as identified by the cluster-based permutation testing. This was only applied to the responses of the training set.

**Figure S3.**
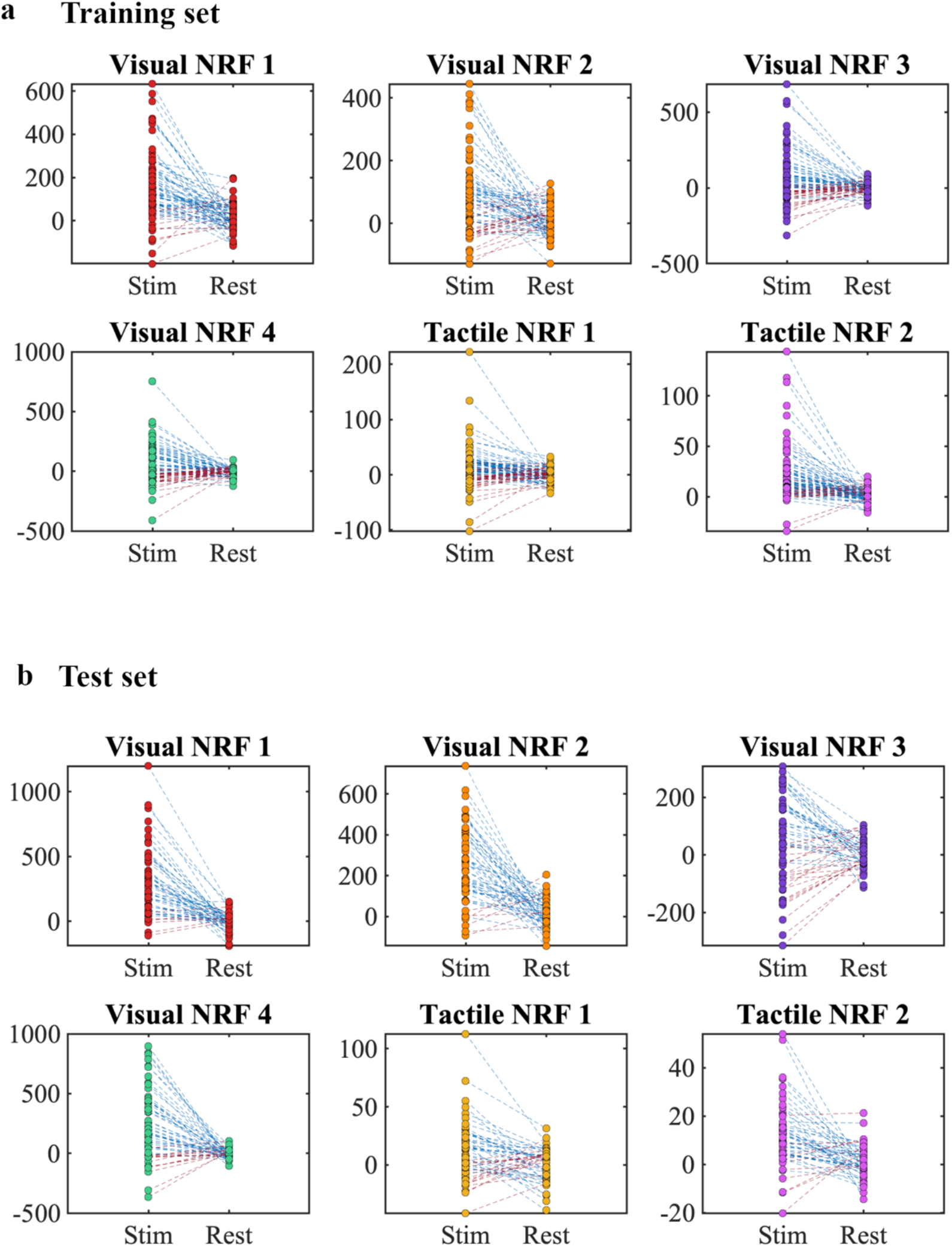
a) Training and **b)** test set magnitudes for stimulus-evoked potentials (Stim) and resting state activity (Rest) using the six neurodynamic response functions (NRFs) identified in the training set. Magnitudes were estimated for the stimulus responses and resting state activity of each recording. Dashed lines connect the two magnitudes of each recording, where blue means a higher magnitude in the stimulus condition relative to the resting state condition and red a lower magnitude for the stimulus condition.

**Figure S4.**
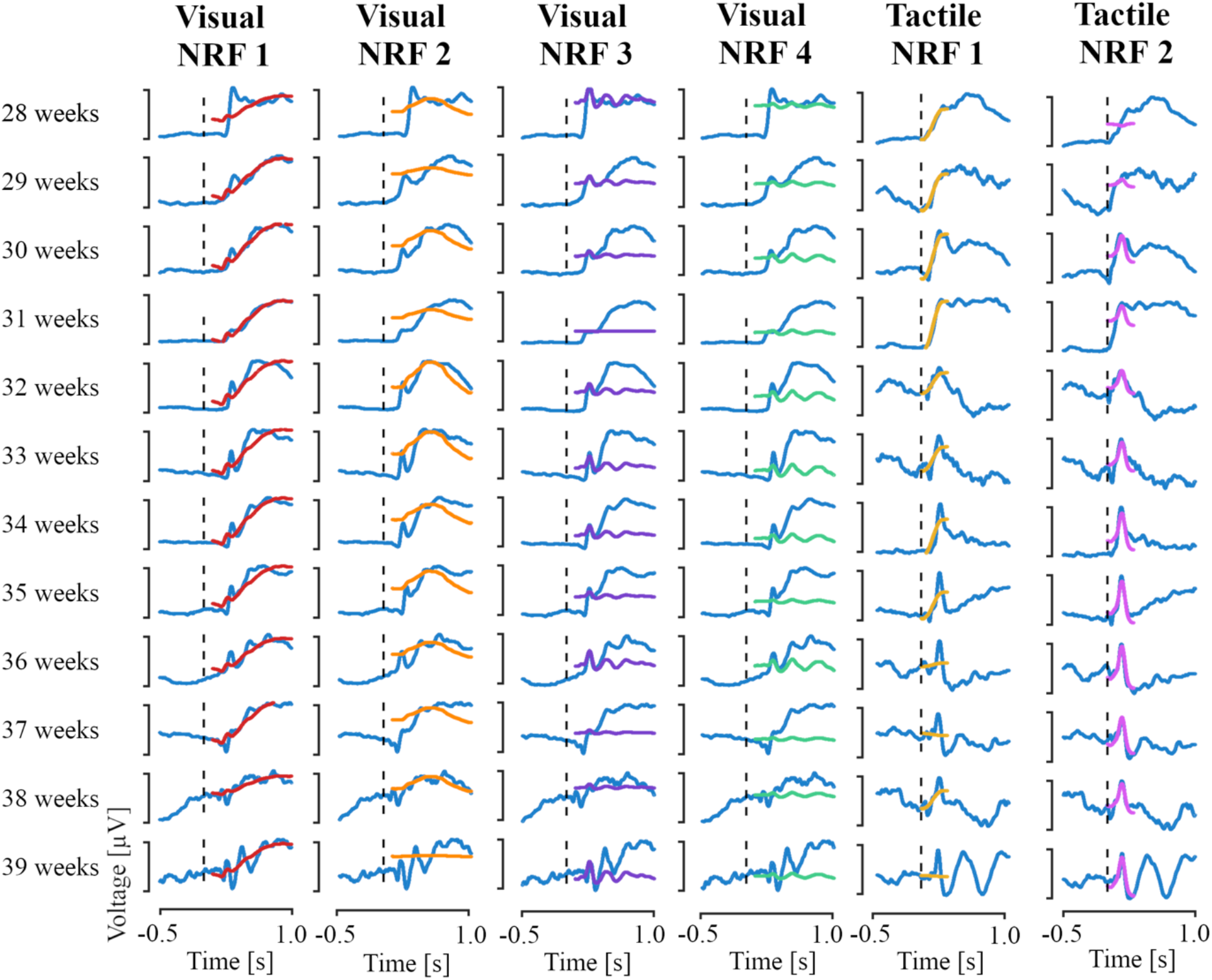
Age-dependent neurodynamic response function (NRF) projections of the visual and tactile NRFs on the training set responses. Each individual projection is plotted on its own individual y-scale. NRFs were projected on age averages after computing age-weighted evoked potentials using linear regression models (see methods). These age-weighted potentials were Woody filtered to the NRF after which the NRF was projected on the EEG traces. Vertical dashed lines correspond to time = 0 seconds (i.e., the stimulus onset).

**Figure S5.**
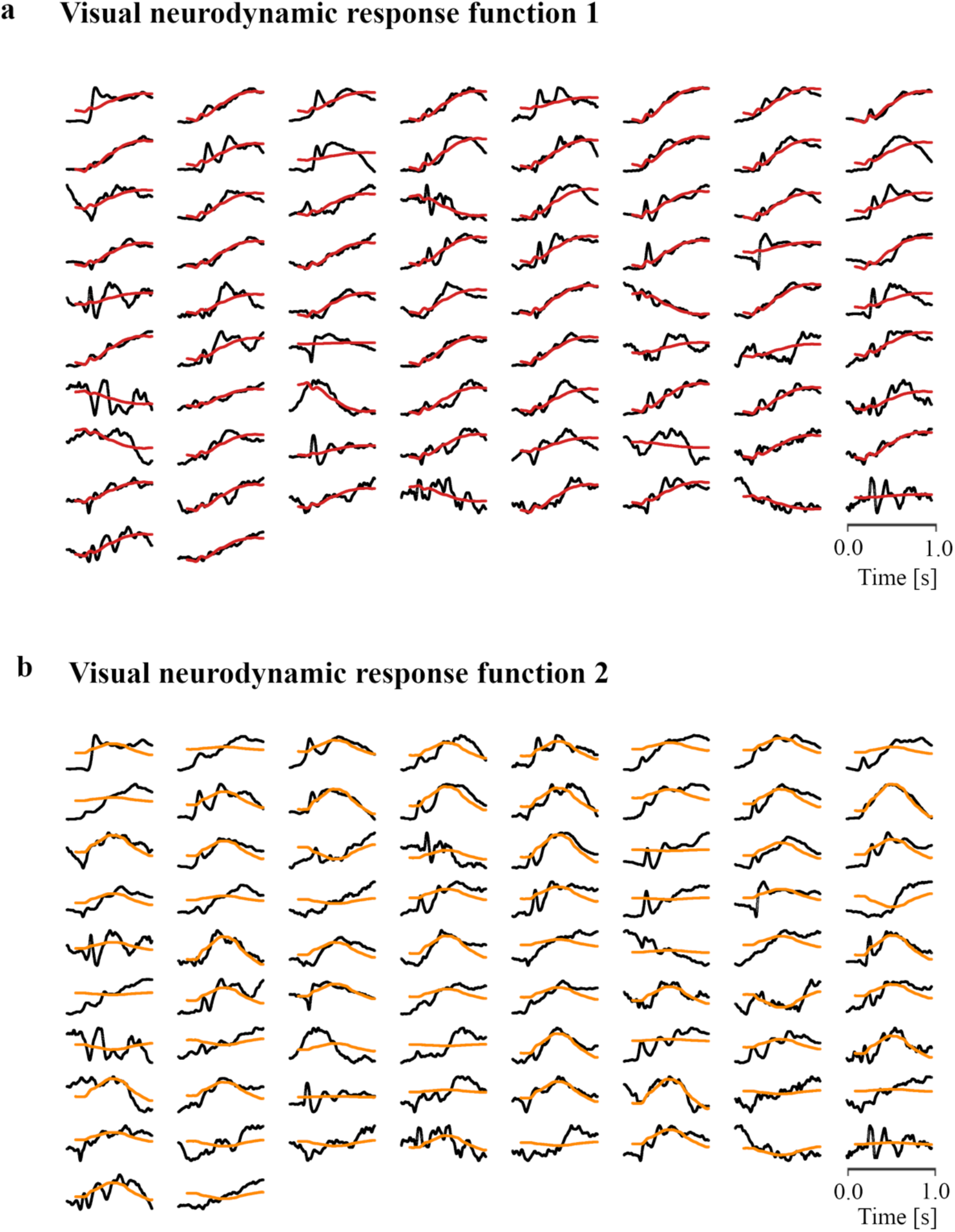
Evoked response function (NRF) projections of visual NRFs 1 and 2 onto each averaged recording stimulus response of the training set. Projections are plotted on their own individual y-scale. Responses are shown in increasing age order, with the youngest subject in the top left.

**Figure S6.**
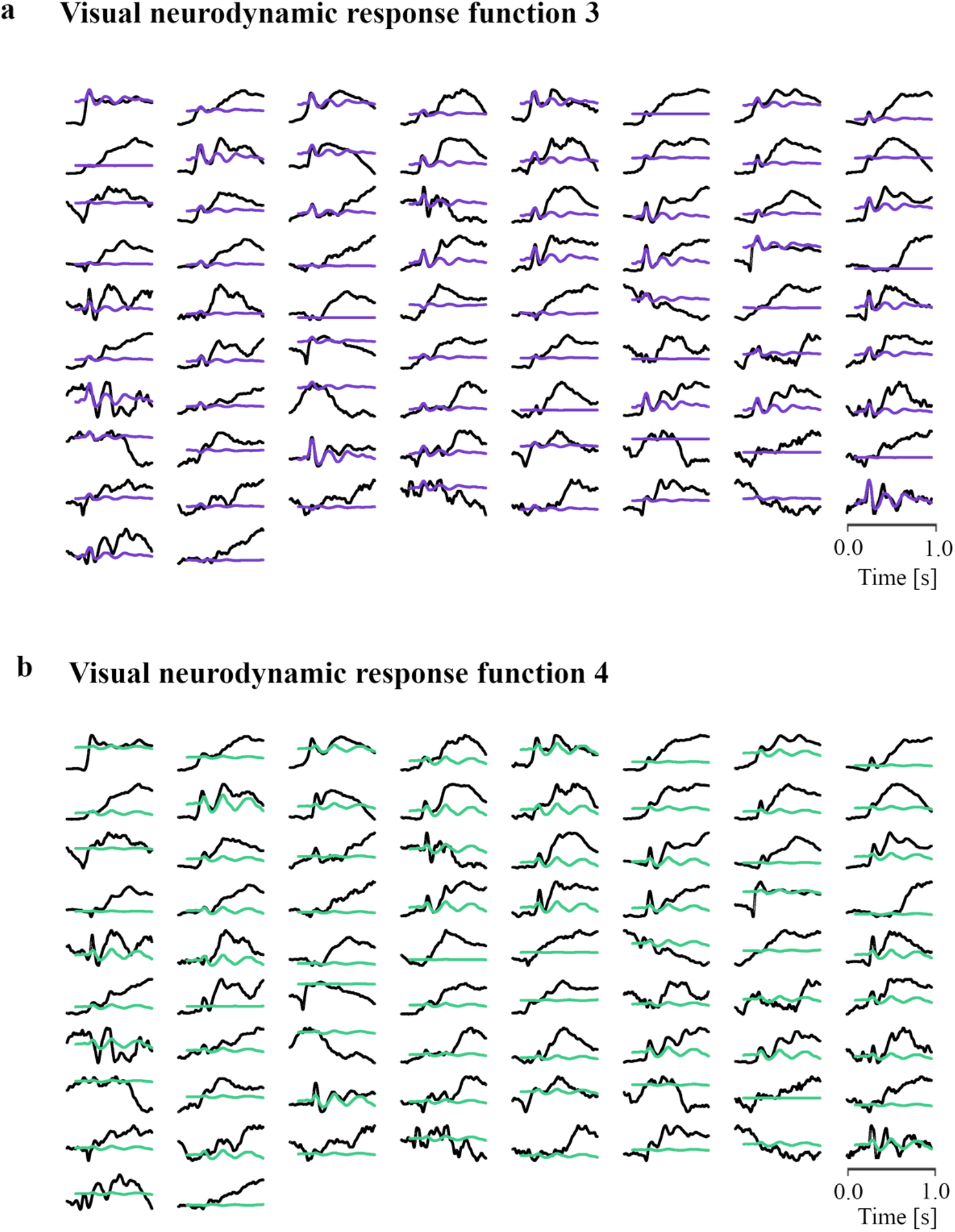
Neurodynamic response function (NRF) projections of visual NRFs 3 and 4 onto each averaged recording stimulus response of the training set. Projections are plotted on their own individual y-scale. Responses are shown in increasing age order, with the youngest subject in the top left.

**Figure S7.**
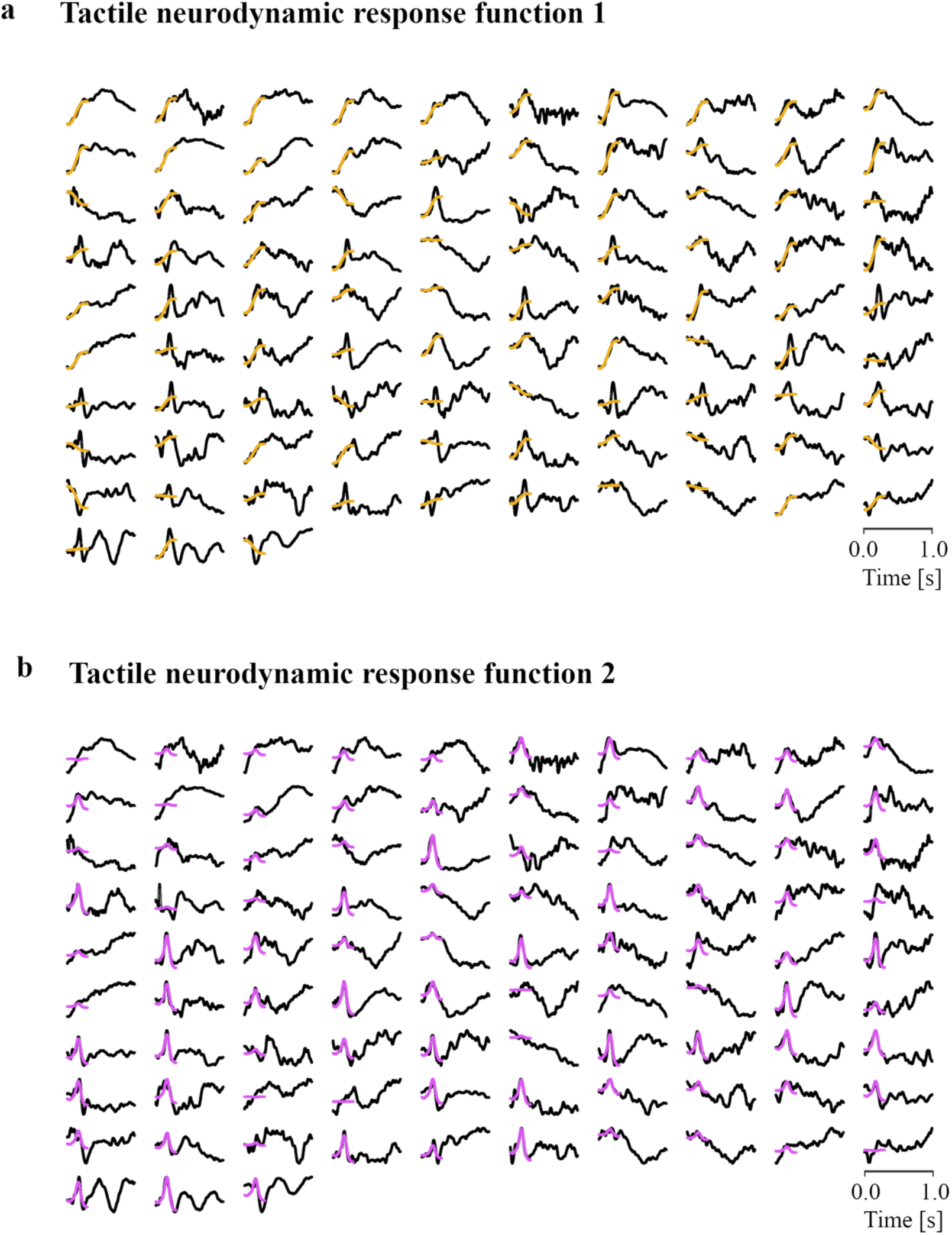
Neurodynamic response function (NRF) projections of the tactile NRFs onto each averaged recording stimulus response of the training set. Projections are plotted on their own individual y-scale. Responses are shown in increasing age order, with the youngest subject in the top left.

**Figure S8.**
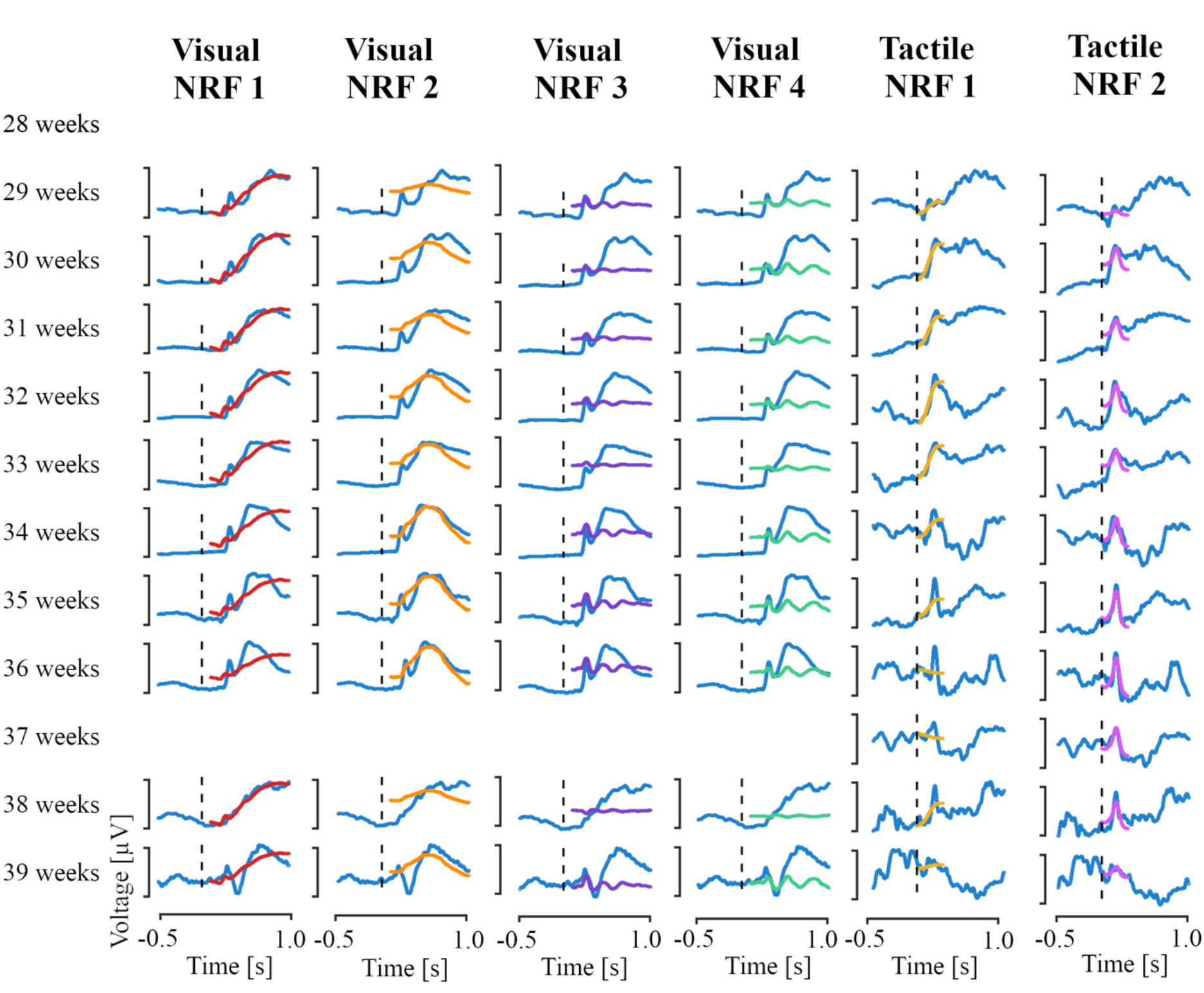
Age-dependent neurodynamic response function (NRF) projections of the visual and tactile NRFs on the test set responses. Each individual projection is plotted on its own individual y-scale. NRFs were projected on age averages after computing age-weighted evoked potentials using linear regression models (see methods). These age-weighted potentials were Woody filtered to the NRF after which the NRF was projected on the potential traces. Vertical dashed lines correspond to time = 0 seconds (i.e., the stimulus onset).

**Figure S9.**
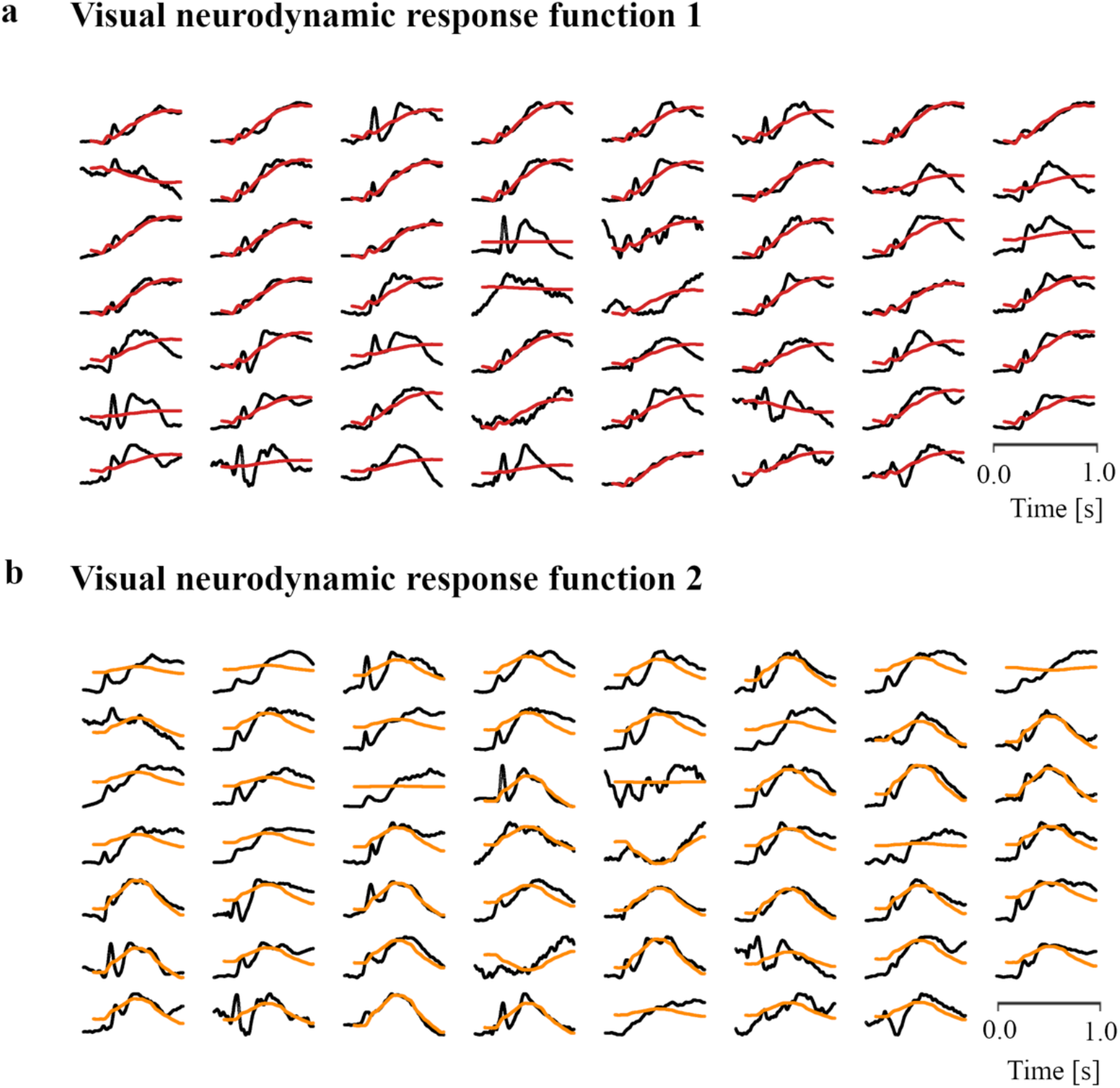
Neurodynamic response function (NRF) projections of the visual NRFs 1 and 2 onto each averaged recording stimulus response of the test set. Projections are plotted on their own individual y-scale. Responses are shown in increasing age order, with the youngest subject in the top left.

**Figure S10.**
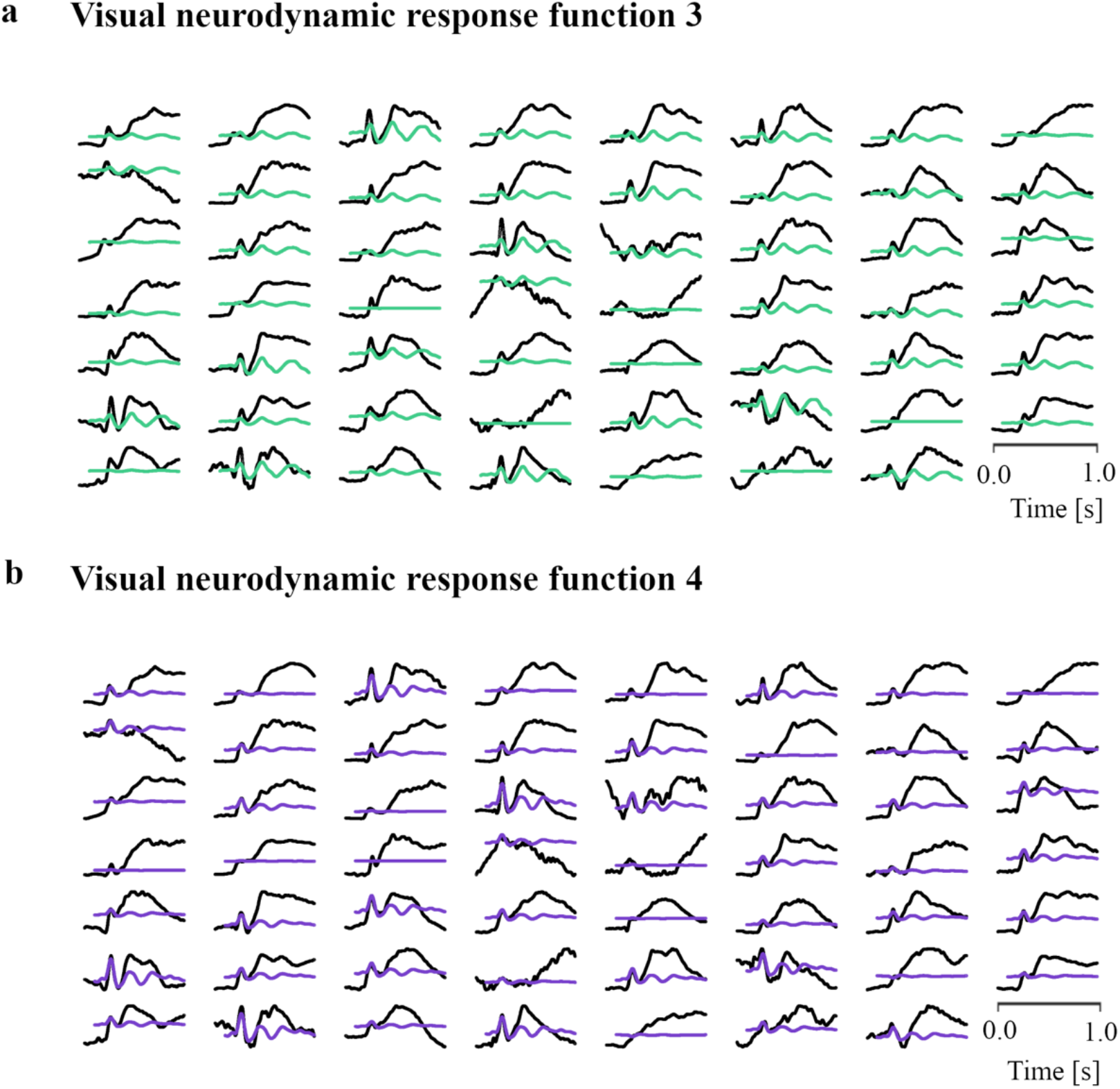
Neurodynamic response function (NRF) projections of the visual NRFs 3 and 4 onto each averaged recording stimulus response of the test set. Projections are plotted on their own individual y-scale. Responses are shown in increasing age order, with the youngest subject in the top left.

**Figure S11.**
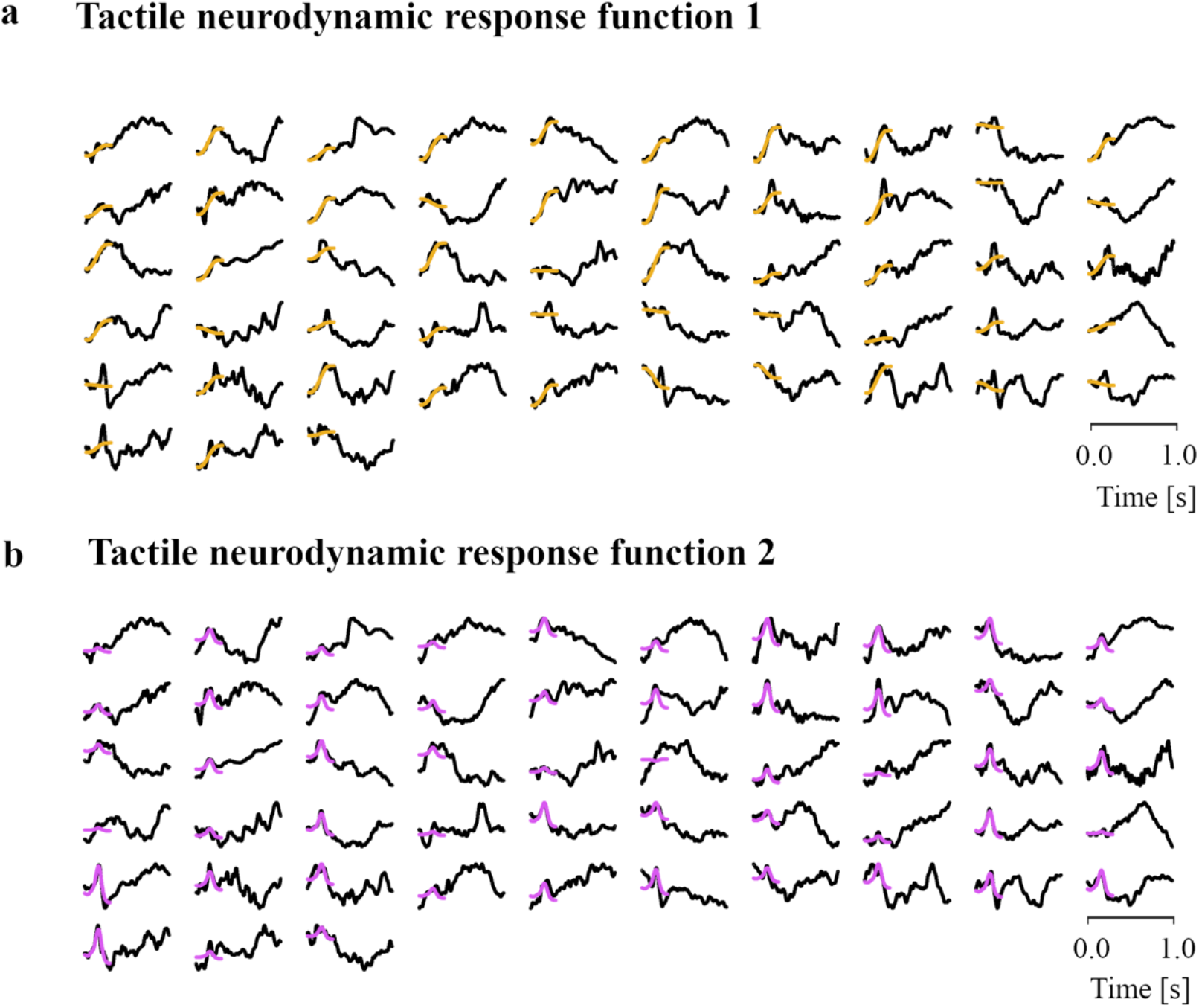
Neurodynamic response function (NRF) projections of the tactile NRFs onto each averaged recording stimulus response of the test set. Projections are plotted on their own individual y-scale. Responses are shown in increasing age order, with the youngest subject in the top left.

**Figure S12.**
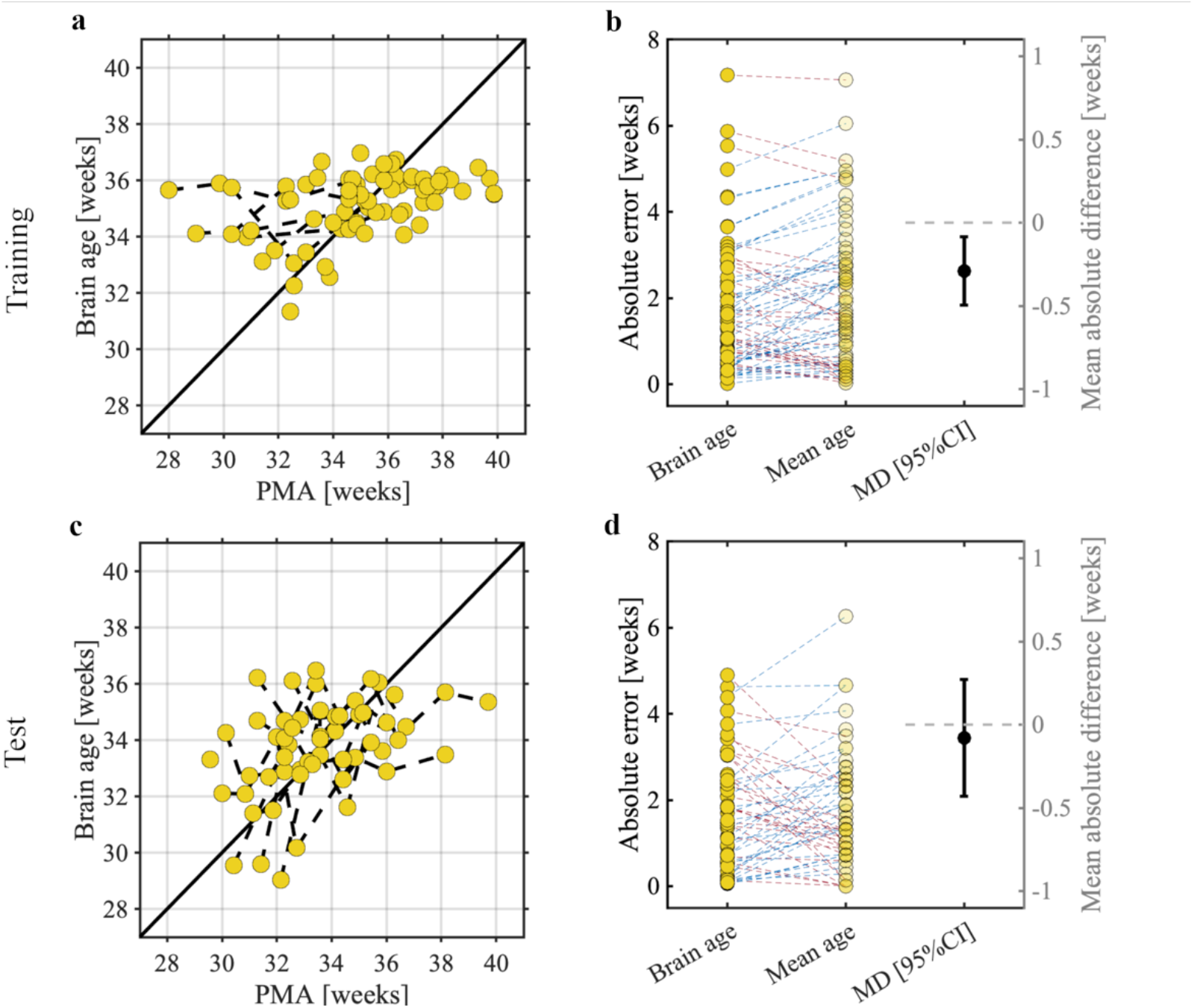
Brain age prediction models and their statistical evaluations for the **a-b)** training and **c-d)** test samples. Panels a and c show the post-menstrual age (PMA) and brain age using leave-one-infant-out cross-validation. Predictions are based on the visual model. Each dot indicates a single recording with PMA predicted using the stimulus responses. Dashed black lines between dots are infants that took part in multiple recordings. Solid black line indicates perfect prediction. Panels b and d depict the comparison in absolute errors between the Brain age and null model (Mean age) and its mean absolute difference including 95% confidence interval (i.e., MD [95%CI]). Blue dashed lines mean a higher absolute error for the mean age prediction relative to the brain age prediction, and red yield a lower absolute error for the mean age.

**Figure S13.**
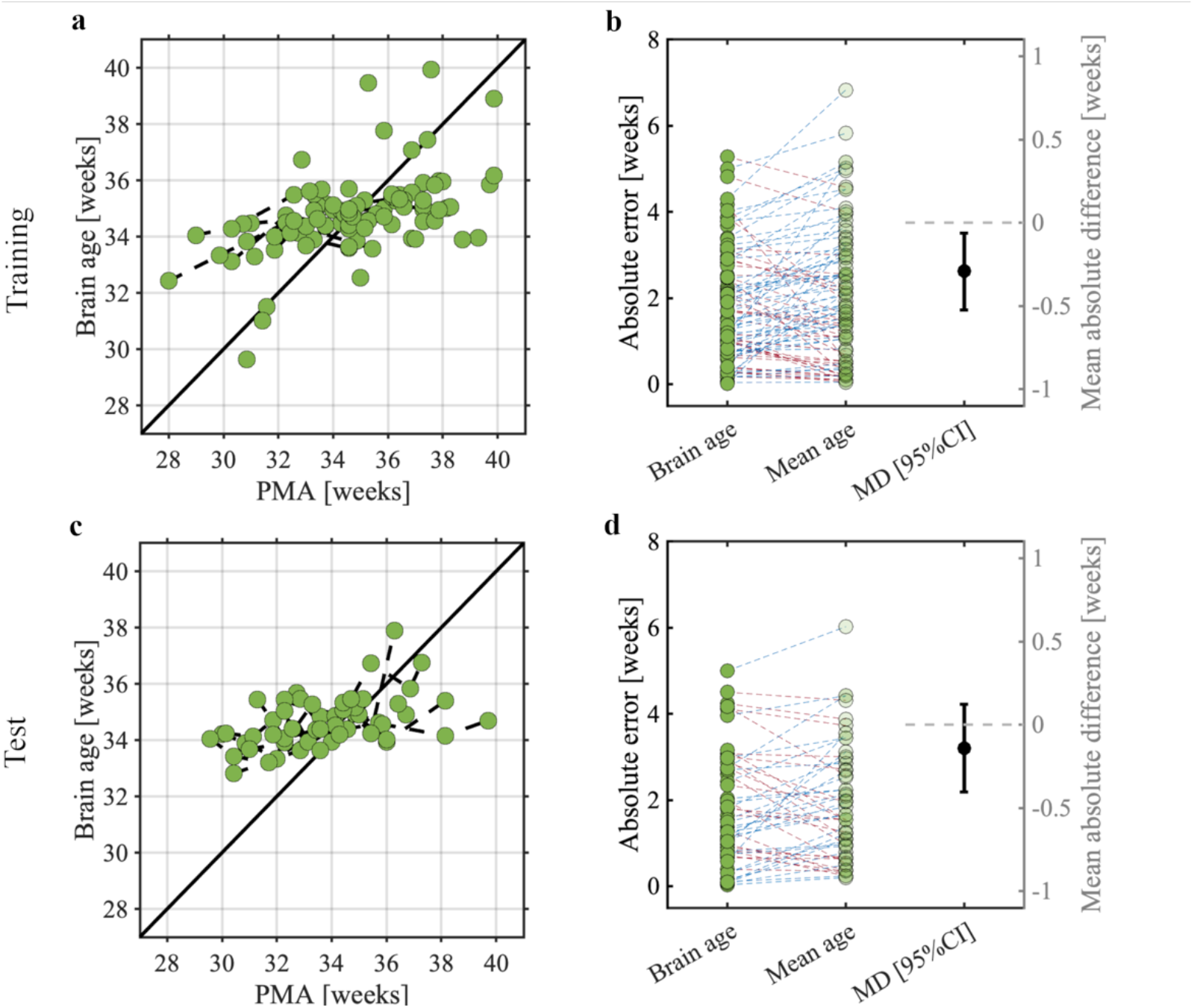
Brain age prediction models and their statistical evaluations for the **a-b)** training and **c-d)** test samples. Panels a and c show the post-menstrual age (PMA) and brain age using leave-one-infant-out cross-validation. Predictions are based on the tactile model. Each dot indicates a single recording with PMA predicted using the stimulus responses. Dashed black lines between dots are infants that took part in multiple recordings. Solid black line indicates perfect prediction. Panels b and d depict the comparison in absolute errors between the Brain age and null model (Mean age) and its mean absolute difference including 95% confidence interval (i.e., MD [95%CI]). Blue dashed lines mean a higher absolute error for the mean age prediction relative to the brain age prediction, and red yield a lower absolute error for the mean age.

**Figure S14.**
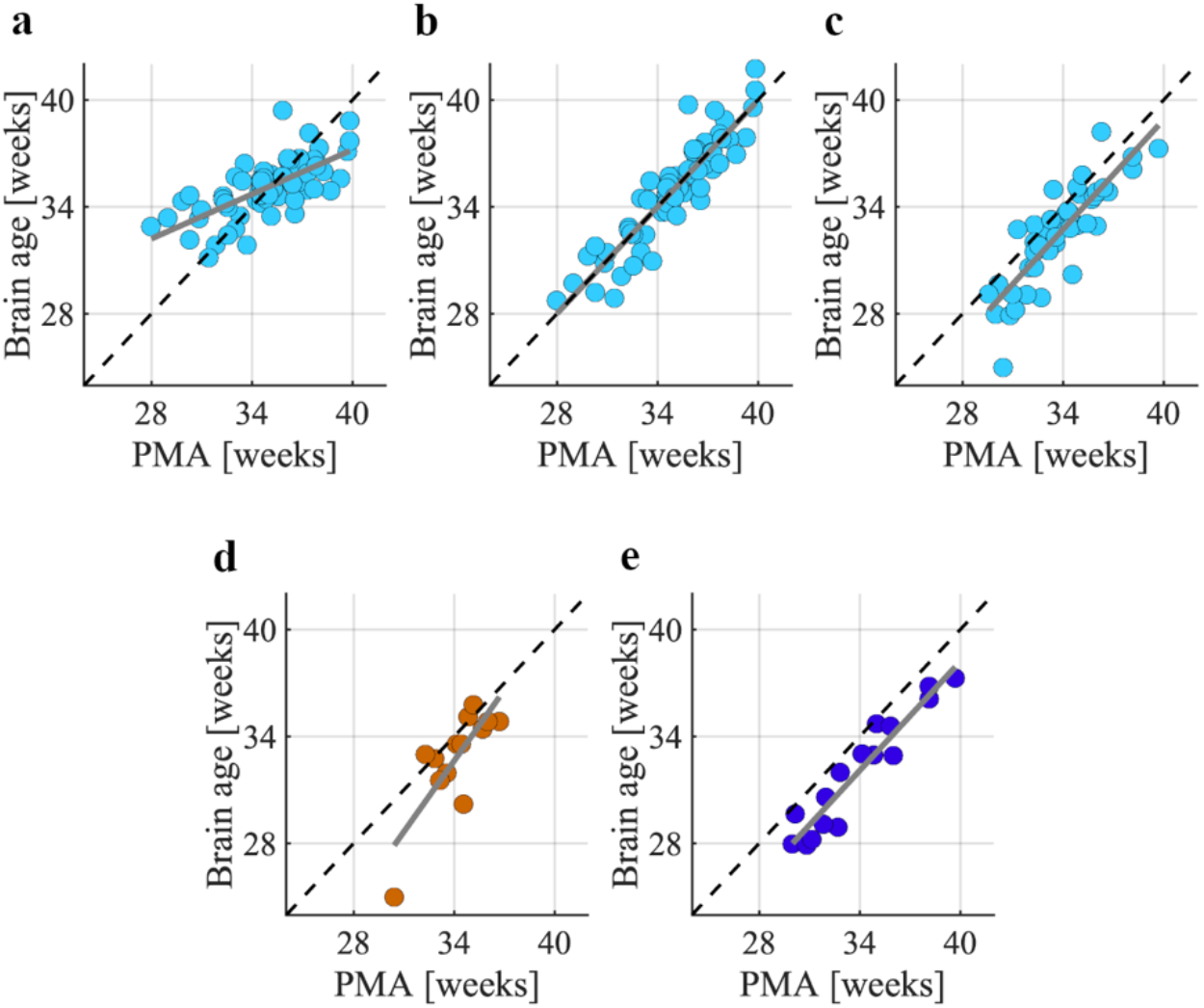
Brain age predictions from which the bias (i.e., deviations from perfect predictions) are removed. Model bias estimated in the **a)** training set by fitting a line of best fit between the brain age and post-menstrual (PMA) data as solid grey graph. Bias was estimated by taking the difference between this line of best fit and the perfect predictions, and subsequently removed from the brain age predictions of the training set, with the resulting brain age predictions shown in panel **b**) Brain age predictions of younger and older babies are particularly adjusted after bias removal. The line of best fit/bias estimated in the training set was used to remove bias in **c)** the test set, **d)** babies in the test set with average Bayley’s outcomes, and **e)** babies in the test set with below average Bayley’s outcomes following bias removal.

**Figure S15.**
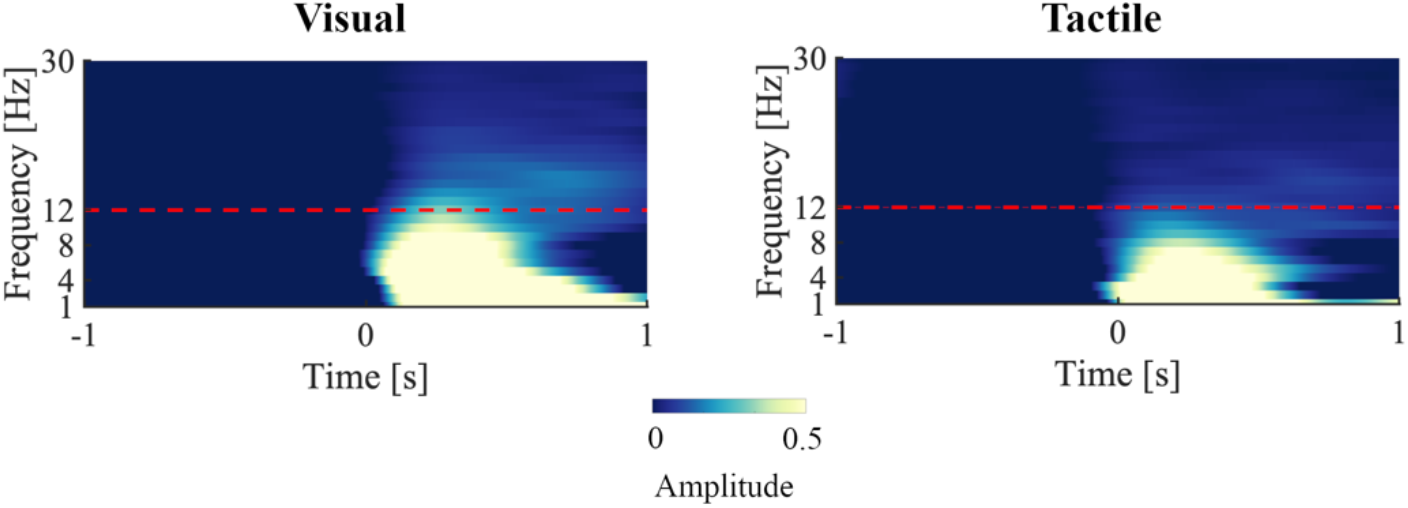
Time-frequency amplitudes of the visual- and tactile-evoked potentials of the training sample. Each time-locked evoked response was bandpass filtered between 1 and 30 Hz with cut-off frequencies of +/-1 Hz around the frequency of interest. Hilbert transforms of the bandpass filtered signals and its instantaneous amplitude was computed by taking the modulus. Time at 0 sec indicates stimulus onset. Horizontal dashed red line corresponds to the upper frequency cut-off used for NRF computation.

